# Visual and Quantitative Analyses of Virus Genomic Sequences using a Metric-based Algorithm

**DOI:** 10.1101/2021.06.17.448868

**Authors:** Alexandra Belinsky, Guennadi A. Kouzaev

## Abstract

This work aims to study the virus RNAs using a novel algorithm for accelerated exploring any-length genomic fragments in sequences using Hamming distance between the binary-expressed characters of an RNA and query patterns. The found repetitive genomic sub-sequences of different lengths were placed on one plot as genomic trajectories (walks) to increase the effectiveness of geometrical multi-scale genomic studies. Primary attention was paid to the building and analysis of the *atg*-triplet walks composing the schemes or skeletons of the viral RNAs. The 1-D distributions of these codon-starting *atg*-triplets were built with the single-symbol walks for full-scale analyses. The visual examination was followed by calculating statistical parameters of genomic sequences, including the estimation of geometry deviation and fractal properties of inter-*atg* distances. This approach was applied to the SARS CoV-2, MERS CoV, Dengue and Ebola viruses, whose complete genomic sequences are taken from GenBank and GISAID databases. The relative stability of these distributions for SARS CoV-2 and MERS CoV viruses was found, unlike the Dengue and Ebola distributions that showed an increased deviation of their geometrical and fractal characteristics of *atg*-distributions. The results of this work can found in classification of the virus families and in the study of their mutation.

## 1. Introduction

A virus is a tiny semi-live unit carrying genetic material (RNA or DNA – double-helix RNA structure) in a protein capsid covered by a lipid coat. The virus penetrates the cell wall and urges this bio-machine to ‘manufacture’ more viruses.

Some viruses are RNA-based and transfer the genetic information by long chains of four organic acids, namely, Adenine (*a*), Cytosine (*c*), Guanine (*g*) and Uracil (*u*) [1]. DNA-based viruses and double-stranded genetic polymers carry the information by four nucleotides, but one of them is Thymine (t) instead of Uracil. In genomic databases, anyway, even the single-stranded viral RNAs are registered as the complementary chains where Thymine substitutes Uracil due to some instrumental specifics [2] that do not hinder the mathematical aspects of the virus theories. These complimentary RNAs will be used further for numerical modelling in our paper.

A complete RNA is a chain of codons (exons) used to transfer genetic information and introns. Unfortunately, the role of the latter is not well known [3]. Sequencing of RNA or DNA is searching and identifying nucleotides by instrumental means. Codons in RNAs start with an ‘*aug*’ combination of nucleotides and end with one of the following three combinations: ‘*uaa*’, ‘*uag*’, or ‘*uga*’.

Some DNA strands consist of billions of nucleotides, so mathematical methods are widely used in genomics [4]. For instance, the RNA symbols are substituted by number values, and this process is called DNA/RNA mapping [5]-[8]. For example, in Ref. [5], eleven methods of numerical representation of genomic sequences are listed and analysed to conclude that each of them is preferable in a particular application, and no universal mapping algorithm is equally advantageous for all genomic study.

Different retrieval algorithms can be applied to genomic sequences, including signal-processing means [7],[9]-[14]. The numerical RNAs can be shown graphically for qualitative analyses. For instance, each nucleotide is represented by a unit vector in a 4-dimensional (4-D) space, and an imaginary walker moves along an RNA sequence, making a trajectory in this space [15],[16]. To avoid apparent difficulties with plotting walks in multi-dimensional spaces, the nucleotides are combined in a certain way [17]. For instance, each nucleotide is associated with one of four unit vectors in 2-D space, which projections on the plane axes can take positive (+1) or negative (–1) values [16],[18]. A trajectory is built moving along the consecutive number of a nucleotide in the studied genomic sequence.

In general, DNA walks allow the detecting of codons and introns, discovering hidden RNA periodicity [12]-[14] and calculating phylogenetic distances between genomic sequences [19], among others. Some additional results and reviews on DNA imaging can be found, for instance, in Refs. [15],[20]-[22], where the necessity to use specified walks for each class of genomic problems is shown.

Genomic walk analysis can be followed by calculating fractal properties of distributions of nucleotides [23]-[32]. Fractals are self-similar or scale-invariant objects. It means that small ‘sub-chain’ geometry is repeated on larger geometry scales, although randomly distorted. A biopolymer chain in a solution is bent in a fractal manner [26]. This fractality influences the chemical reaction rate, diffusion, and surface absorption of long-chain and globular molecules, among others [25],[26],[33]-[36]. Because polar solvents have frequency-dependent properties, they are adjusted by applied microwaves that influence the polymer fractal dimension. Thus, some bioreactions can be controlled by a weak high-gradient microwave field [37],[38].

Although many achievements are known in the numerical mapping of RNAs, some questions have not been resolved. For instance, the known genomic walks are designed to track single nucleotides or their pairs, leading to crowdy trajectories and overloaded plots that are challenging to analyse visually in one plot [5]-[8]. Meanwhile, the complete RNAs of viruses are composed mostly of codons, and one repetitive pattern therein is *atg-*triplet. We propose that these triplets build the viral RNA scheme or skeleton.

Our developed pattern search algorithm calculates the triplet distributions along an RNA sequence. Additionally, the same algorithm can make the walks of each of the four nucleotides. These trajectories, found not woved strongly in comparison to [5],[8] and being imaged by different graphical means on the same figure and equipped with interactive links to the names of genes, make visual analyses much more effective. The results of the creation of such a tool and applications to genomes of several viruses are given here. These codes and graphic means can be applied for research and practical applications. One of them is the studies of the stability of mentioned *atg*-schemes towards mutations, variation of codon fillings and the fractality of *atg*-distributions, among others.

In Section 2, the developed calculation algorithms and plotting techniques are considered in detail. The results of using these techniques to the SARS CoV-2, MERS CoV, Dengue and Ebola viruses are in Section 3. They are discussed in Section 4, and conclusions are rendered in Section 5. The text is followed by a list of more than 70 references. In Appendix 1, all necessary data for the analysed virus RNAs taken from GenBank and GISAID are given in a tabular form, including the parameters calculated in this contribution.

## 2. Materials and Methods

### 2.1. Materials and Data Availability

In this paper, the arbitrary-chosen complete genomic sequences are studied taken from GenBank [39] and GISAID [40]. Among them, 21 SARS CoV-2 genomic sequences from GISAID and one from GenBank, ten genomic sequences for the MERS coronavirus (GenBank), 25 genomic sequences for the Dengue virus from GenBank and 15 Ebola virus genomic sequences (GenBank). Data from the GISAID database are available after registration. All names of genomic sequences are given in figure legends and in Tables 1–4.

### 2.2. Methods

#### 2.2.1. Metric-based atg-Triplet Walking Algorithm

As it has been stated above, for both DNA and RNA descriptions by characters, their alphabet consists of four nucleotides. These designations are used to study RNA and DNA if their physicochemical properties are outside the research scope.

In many cases, the RNAs/DNAs have repeated patterns of nucleotide sequences, and these regions are better conserved in mutations [9]. Systems of these repeating fragments are considered as skeletons or schemes of these chains [41],[42]. As a rule, pattern discovery relates to nondeterministic polynomial time problems (NP-problems), i.e. solution time increases exponentially with the sequence length.

A typical algorithm compares a query character pattern with a length of nucleotides with the following one-symbol shift of query along a chain. In our code, we use these techniques, as well. Usually, the search algorithms working with characters having no assigned numerical values, are slower in 1.43-2.37 times than ones processing the binary variables [43],[44]. Then, in our code, all RNA’s characters are transformed into binaries before all operations using a Matlab operator dec2bin(*character*) [45]. In computers, for instance, the UTF-8 format allows encoding all 1,112,064 valid character code points, and it is widely used for the World Wide Web [46]. The first 128 characters (US-ASCII) require only one byte (eight binary numbers) in this format. If binary units initially represent the DNA sequences, then calculating the DNA sequence’s numerical properties in this form reduces the computation time.

Because binary sequences now write the DNA/RNA chains, they can be characterised quantitatively using a suitable technique. One calculates a metric distance between the binary-represented symbols and a query (base) ‘moving’ along a chain. Many metric types are used in codes and big data [47]-[55]. The advantage of using metric estimates is that they can be applied in cluster analysis for similar grouping nucleotide or protein distribution patterns. For instance, it can help to class the virus RNAs [54]. Particularly, this distance can be the Hamming one [47],[48] used further in this paper.

Consider a flow chart of our code on exploring patterns of arbitrary length *n* (Fig. 1). It starts with importing sequence data *A* of the length *N* from any genomic database in FASTA format [39],[40] and defining a query pattern *B* of any length *n*. From both files, the empty spots should be removed [56]. In the second step, the data files are transformed into binary strings, and they are used to calculate Hamming distance between each binary symbol of *A* and *B*. This distance is a metric for comparing two binary values, and it is the number of bit positions in which the two bits are different. To calculate Hamming distance *d_H_* between two strings *A* and *B*, the XOR operation (*A* ⊕ *B*) is used, and the total number of ‘1’s in the resultant string *C* is counted. If the compared binary-represented symbols are the same, the distance value is zero. Only *n* characters are compared on each count; then, the query is moved on one symbol, and calculations are repeated. Then, a string *C* of numerical estimates of the length *M* = *n* × *N* is a product of step 3 of this code. Not all registered genomic sequences are divisible by *n*. In this case, a needed number of characters *a* is added to the end of the sequence *A*, or the Hamming metric operation can be fulfilled using Levenstein’s distance formula from Ref. [50], workable for compared sub-strings of arbitrary length [53].

**Fig. 1.**
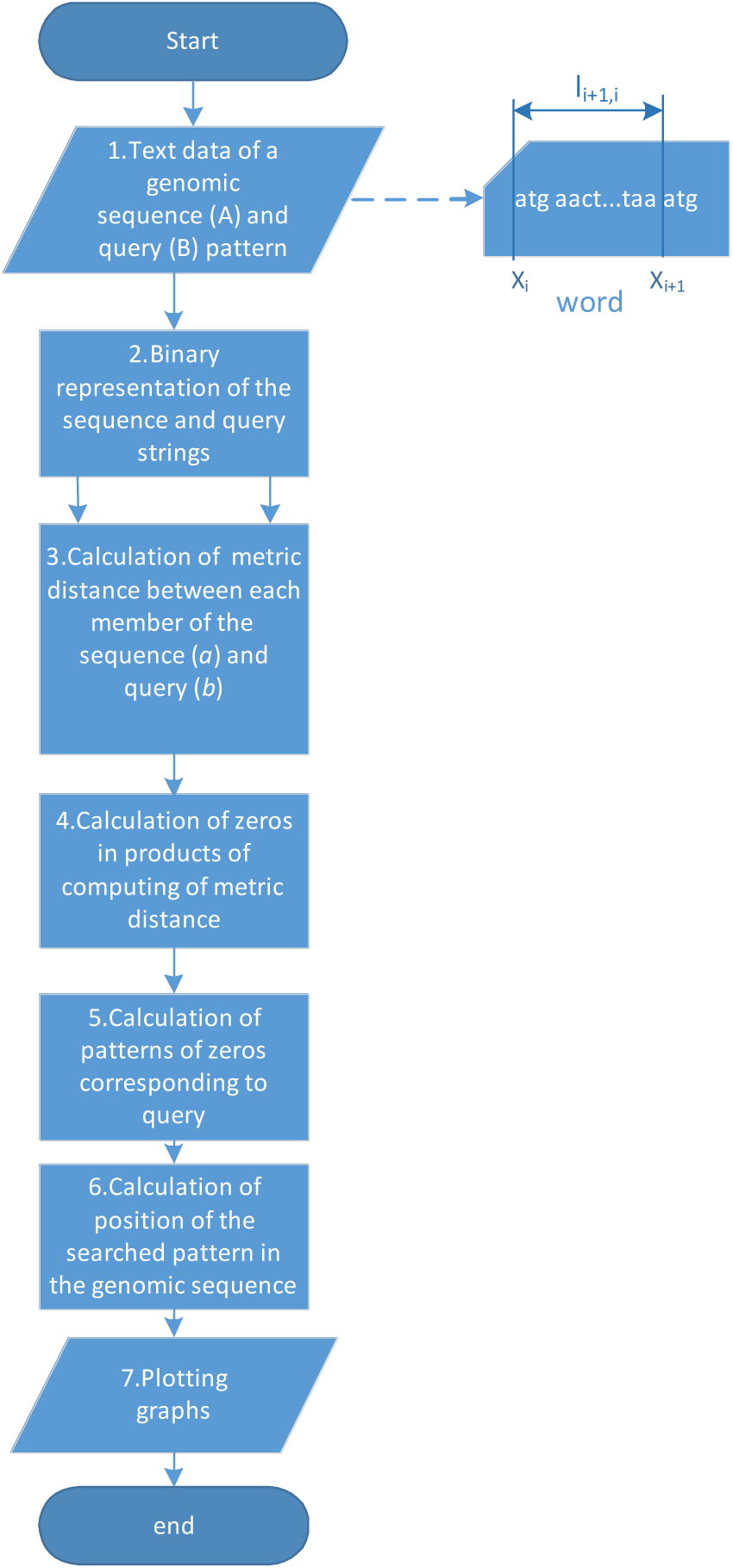
Algorithm flowchart.

The following two parts of our algorithm are with calculation of numbered (*y_i_*) query positions in the RNA sequence *x_i_* according to the Hamming-distance data. All zeros in the string *C* are initially obtained (step 4, Fig. 1). Then, only *n* neighbouring zeros corresponding to a query are selected (step 5, Fig. 1), and this query is numbered in a sequential manner starting with the first one found in RNA. The positions *x_i_* of these numbered queries *y_i_* in a complete RNA sequence are calculated analytically (step 6, Fig. 1). Let us take the coordinate of the first symbol in a numbered found query, then a set of points can be built along a studied sequence. These points, being connected, make a curve called a query walk.

In this paper, the start-up *atg*-triplet is used below as a query (pattern *B*). We define the positions *x_i_* of the first symbols of the sequentially numbered *atg*-triplets in an RNA *A*, and the *atg*-walks are plotted. Additionally, we calculate the word length 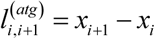. In our algorithm, a ‘word’ is a nucleotide sequence starting with an *atg*-triplet and all symbols up to the next *atg*-one (Fig. 1, right side).

The proposed algorithm was realised in the Matlab environment [45], and it is a few-ten-line code. The following Matlab library functions were used:

1. *dec*2*bin*(character variable) *–* to transform a character variable into a binary one
2. *ptisd* 2(*a*,*b*, ‘*hamming*’) values *a* and *b –* to calculate the Hamming distance value between two binary
3. *zeros* (string) *–* calculation of numbers of zero-values in a string
4. *plot* (*y*, *x*) – plot line function *y* (*x*)

The developed algorithm was applied to many available virus complete genomes to validate it, and the calculated *atg*-positions were compared with those available from databases.

#### 2.2.2. Visualisation Techniques

##### 2.2.2.1. The *atg*-Walks

The viral RNAs, consisting of thousands of nucleotides, are challenging to analyse, and many visualising methods are used. Among them are plotting the DNA walks projected on the spaces of appropriated dimensions considered above. Some contributions are full of symbolic designations of nucleotides and diagrams showing positions of genes in the complete DNA sequences, among others. The preference for a visualisation method is dictated by the specificity of applications, although there is a need for a universal graphical tool.

This paper found that *atg*-walks could be plotted by coordinates of the numbered *atg*-triplets in a complete RNA sequence. Like the known DNA walks, these dots can be considered the points of a trajectory named here as *atg*-walk (Fig. 2A). A diagram showing positions of the symbol *a* in defined *atg-*triplets is given in Fig. 2B.

**Fig. 2.**
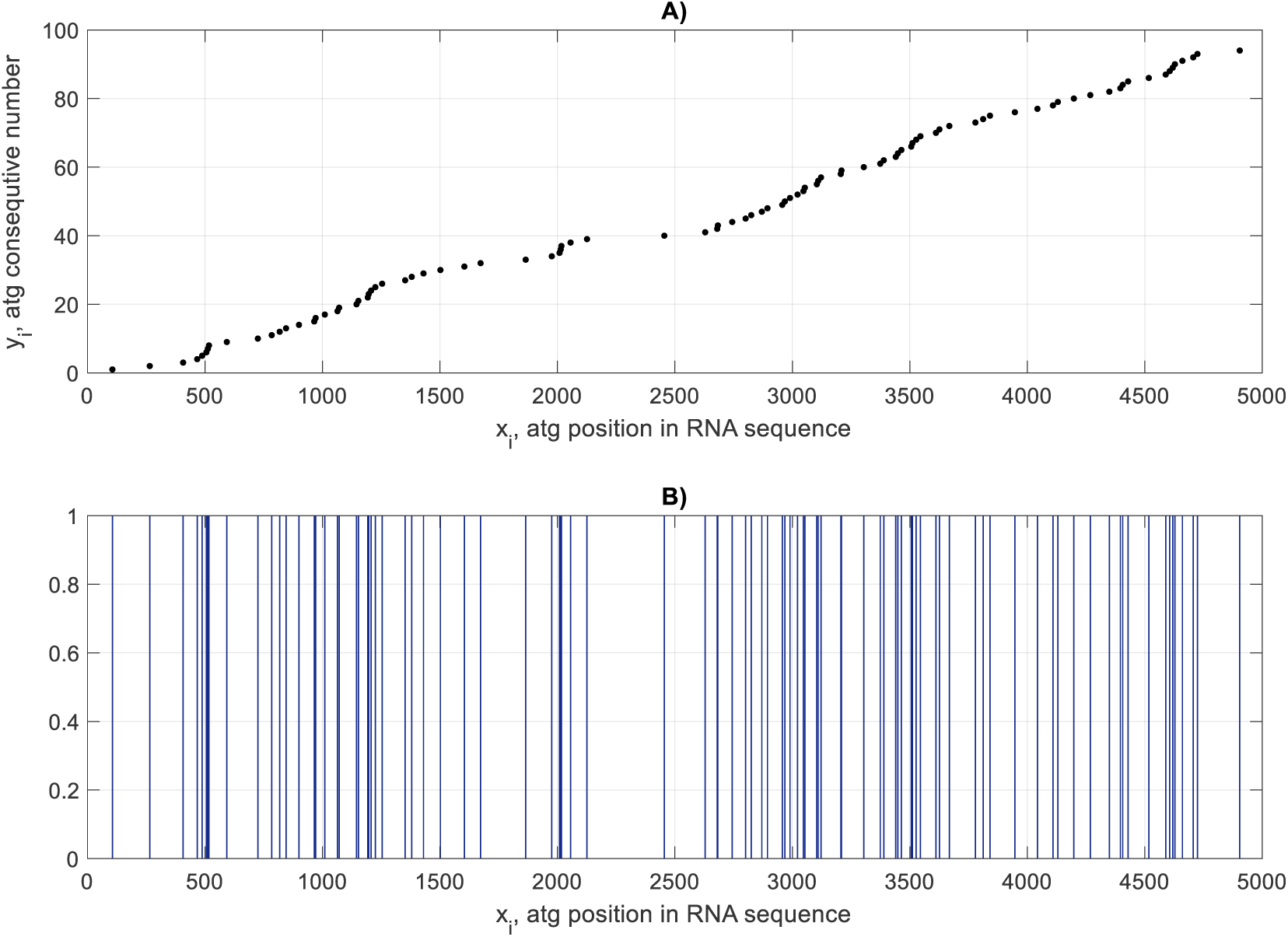
Positions of *atg*-triplets along the genome sequence of SARS-CoV-2 virus MN988668.1 (GenBank) given for the first 5 000 nucleotides provided by points (A) and vertical blue lines in diagram (B)

Like known single-symbol DNA walks, the *atg*-trajectories have fractal properties. Their type was defined by analysing distributions of the coordinates of *atg*-triplets shown by vertical lines along an RNA sequence (Fig. 2B). These *atg*-distributions have repeating motifs on different geometry scale levels, i.e. they can have fractal properties. Below, our initial assumption about the fractality of *atg*-distributions is confirmed: we calculated the fractal dimensions of complete genomes of several tens of virus sequences. Presumably, the *atg*-triplets are distributed along with the RNA sequences of studied viruses according to the random Cantor multifractal law [51].

##### 2.2.2.2. Multi-scale Mapping of RNA Sequences

It is necessary to see full-scale virus RNA maps and analyse all types of mutations. Previously, the most attention was paid to mapping *atg*-triplets, thinking that they constructed a skeleton of RNA, a relatively stable structure. Besides the structural mutations changing the *atg*-distributions, the nucleotides vary their positions inside codons. Our algorithm considers even a single symbol as a pattern, and it allows the calculation of distribution curves for each nucleotide similarly to *atg*-ones. These curves can be considered as the first level of spatial detailing of RNAs. The words in our definitions (see Fig. 1) compose the second level. They compose a gene responsible for synthesising several proteins, and the genes belong to the third level of detailed visualization of RNAs.

A combined plotting of elements of the hierarchical RNAs organisation will be helpful in the visual analyses of RNAs and DNAs. One of the ways is shown in Fig. 3. Here, positions of *a*-symbols of *atg*-triplets in an RNA sequence are given by vertical blue lines (second hierarchical level of visualization). Words take spaces between these vertical lines (see Fig. 1). They are filled by numbered nucleotides (points of a different colour), which are the first level of RNA detailed imaging. This allows distinguishing nucleotides even at the beginning of coordinates where crowdedness is seen (inlet in Fig. 3).

**Fig. 3.**
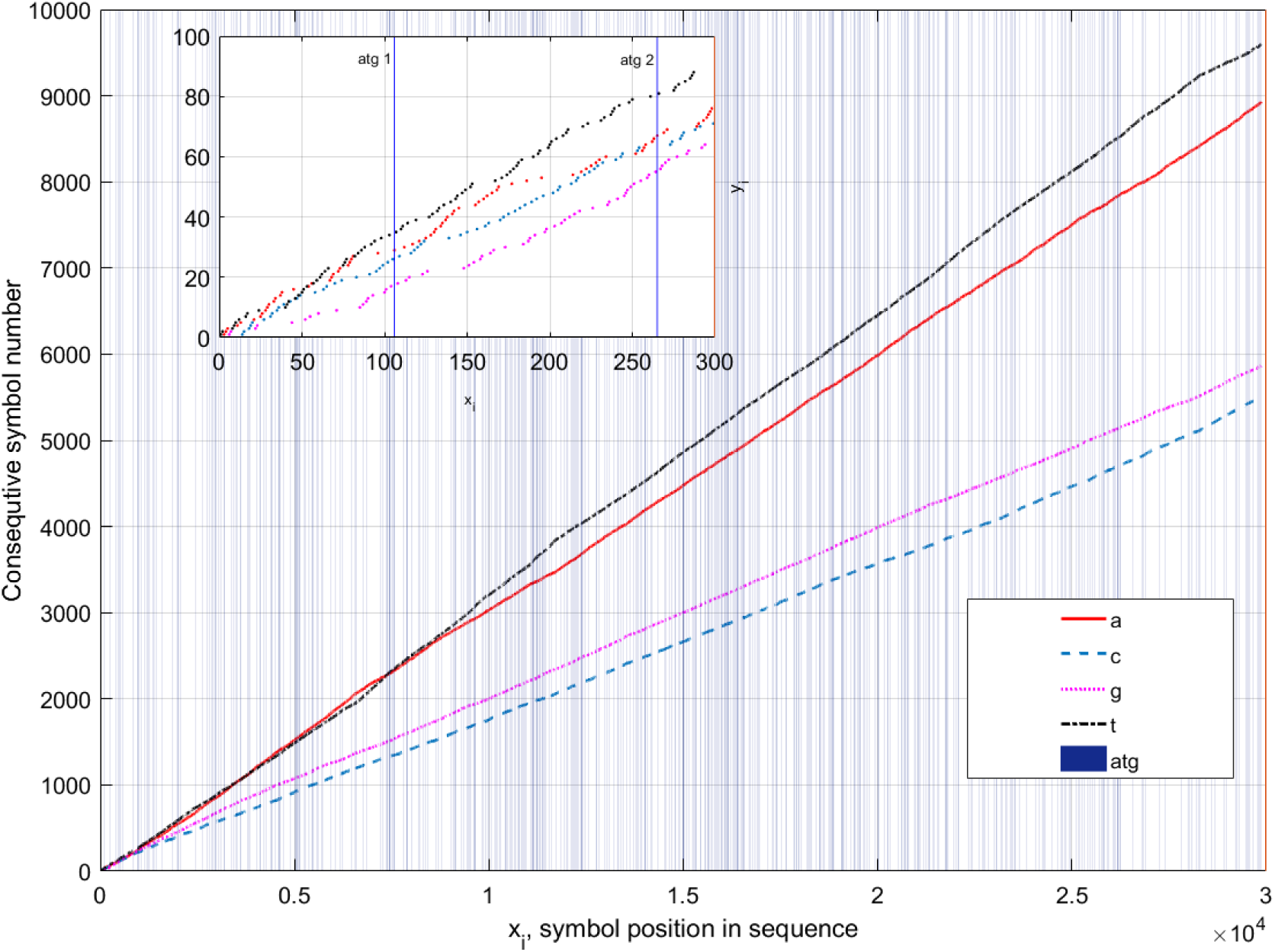
Two-scale study results of a SARS-CoV-2 virus MN988668.1 (GenBank). In the inlet, these symbols are pointed inside the words given for the first 300 nucleotides. Grid lines are in black colour.

The next level of hierarchical RNAs organisation is with genes. For instance, in GenBank [39], the list of symbols of an RNA sequence in FASTA format is followed by a diagram where the genes are given by horizontal bars with the gene’s literal designations. In our case, this diagram can be attached to a two-scale plot considered above. Another solution is to equip our figures with gene hyperlinks, an interactive means highlighting whose genes a nucleotide or codon belongs to, shown by a pointer. This application for third level visualisation is currently under development.

Thus, the developed pattern search algorithm based on the use of Hamming distance applied to binary representations of nucleotide symbols allows building combined plotting of hierarchical organisation of the RNAs of viruses. It can also be applied to the analyses of more complex protein structures.

#### 2.2.3. Calculation of Fractal Dimension of atg-Triplet Distributions

In many previous studies, the fractality of distributions of nucleotides along with the DNA/RNA sequences has been studied [6],[16]-[22],[23],[24],[27]-[36],[51]-[60]. The motifs of small-size patterns are repeated on large-scale levels. Thus, the nucleotide distribution along a genome is not entirely random due to this long-range fractal correlation of amino acids, as is mentioned in many papers. The measure of self-similarity is its fractal dimension *d _F_* approaches. that can be calculated using different

The large-size genomic data are often patterned, and each pattern can have its fractal dimension, i.e., the sequences can be multifractals [31]. This effect is typical in genomics, but it is also common in the theory of nonlinear dynamical systems, signal processing and brain tissue morphology, among others [51]-[63].

Discovering the fractality of genomic sequences is preceded by their numerical representation, for instance, by walks of different types [20],[16],[6]-[8],[27],[31]. Then, each step value of a chosen walk is considered a sample of a continuous function, and the methods of signal processing theory are applied [12],[13].

In our case, the fractal dimension calculations can be applied directly to the distribution of *atg*-consequitive numbers *y_i_* (Fig. 2a), similarly to the cited above works. Unfortunately, in our case, considering about straight-line distribution of *atg*-triplets *y_i_* (*x_i_*), the fractal dimension of these curves is close to one, and it is insensitive against many RNA variations. Instead, the codon’s word-length 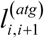 distributions (see Fig. 1, right part) along the RNAs sequences are proposed to use.

A particular distribution of the word lengths is shown in Fig. 4 by bars whose heights equal the word lengths. Then, the algorithms, usually applied to the sampled signals, can be used to compute the statistical properties of these word-length distributions.

**Fig. 4.**
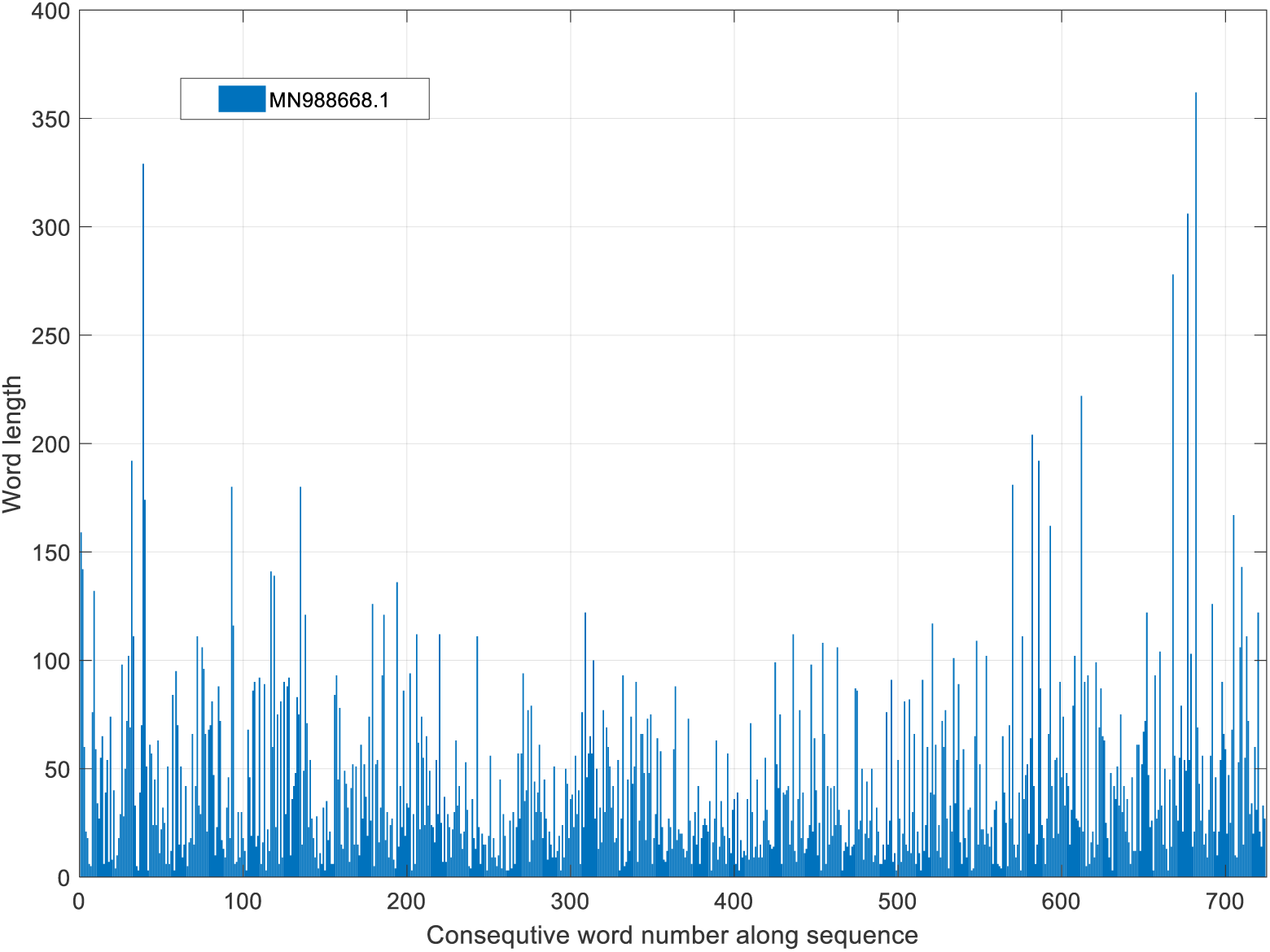
Word-length 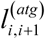 distribution in a SARS Cov-2 virus MN988668.1 sequence (row 1, Table 1, Appendix 1).

In this paper, the fractal dimension of word lenghth distributions and distribution of *y_i_* was calculated using a software package FracLab 2.2 [64]. Although many researchers tested this code, it is again verified to calculate the Weierstrass function, which is synthesised according to a given value of the fractal dimension [65]. This code provides results with reasonable accuracy if the default parameters of FracLab are used.

In a strong sense, the fractal dimension was defined for the infinite sequences. In our case, the studied RNAs have only 268-730 *atg*-triplets depending on the virus. Then, the fractal dimension values were estimated approximately. This is acceptable for our analysis of even short-length RNA sequence viruses like Ebola (Table 4).

## 3. Results

### 3.1. Study of *atg*-Walks of the Complete Genome Sequences of the SARS CoV-2 Virus

In this study, essential attention was paid to studying SARS CoV-2 complete RNA genome sequences. A recent comprehensive review on the genomics of this virus can be found in Refs. [66],[67], for instance. The data used here and throughout this whole paper are from two genetic databases: GenBank [39] and GISAID [40]. A part of the studied genome sequences for this and other viruses is provided in Appendix 1.

Here, the main unit, called a ‘word’, is a nucleotide sequence starting with ‘*atg*’ and the symbols up to the next starting triplet (Fig. 1). The number of *atg*s was calculated by our code and verified by a Matlab function *count* (*A*, ‘*atg* ‘). These results are shown in the third columns, Tables 1–4 (See Appendix 1). The Matlab functions *median*(*L*_word_) and *rms* (*L*_word_) calculated the median and root-mean-square (R.M.S) values of each sequence’s word-length *l*^(^*^atg^*) distribution, correspondingly. The results are in columns 4 and 5 of the mentioned tables.

Consider applying the developed approach to the complete genome of a Wuhan RNA sample MN988668.1 (GenBank) as an example. It consists of 29881 nucleotides and 725 *atg*-triplets (See row 1, Table 1, Appendix 1). Fig. 2 shows the distribution by points of *atg*-triplets for the first 5000-nucleotides of this complete genome.

Fig. 5 illustrates the distribution (in lines) of *atg*-triplets along with complete genome sequences for twenty-one SARS CoV-2 viruses, including Delta, Omicron and a bat-corona sample taken from GenBank [39] and GISAID [40] databases (see Table 1, Appendix 1). There were relatively compact localisations of triplet curves despite the viruses being of different clades and lines. For instance, the divergence of these curves estimated at *x_i_* = 29000 is around only 1%. This confirms the conclusions of many specialists that no new recombined strains have appeared up to this moment, despite many mutations found to date, including the last Omicron lineage [68]. It means that the virus is rather resistant or stable towards forming new families with distinctive properties.

**Fig. 5.**
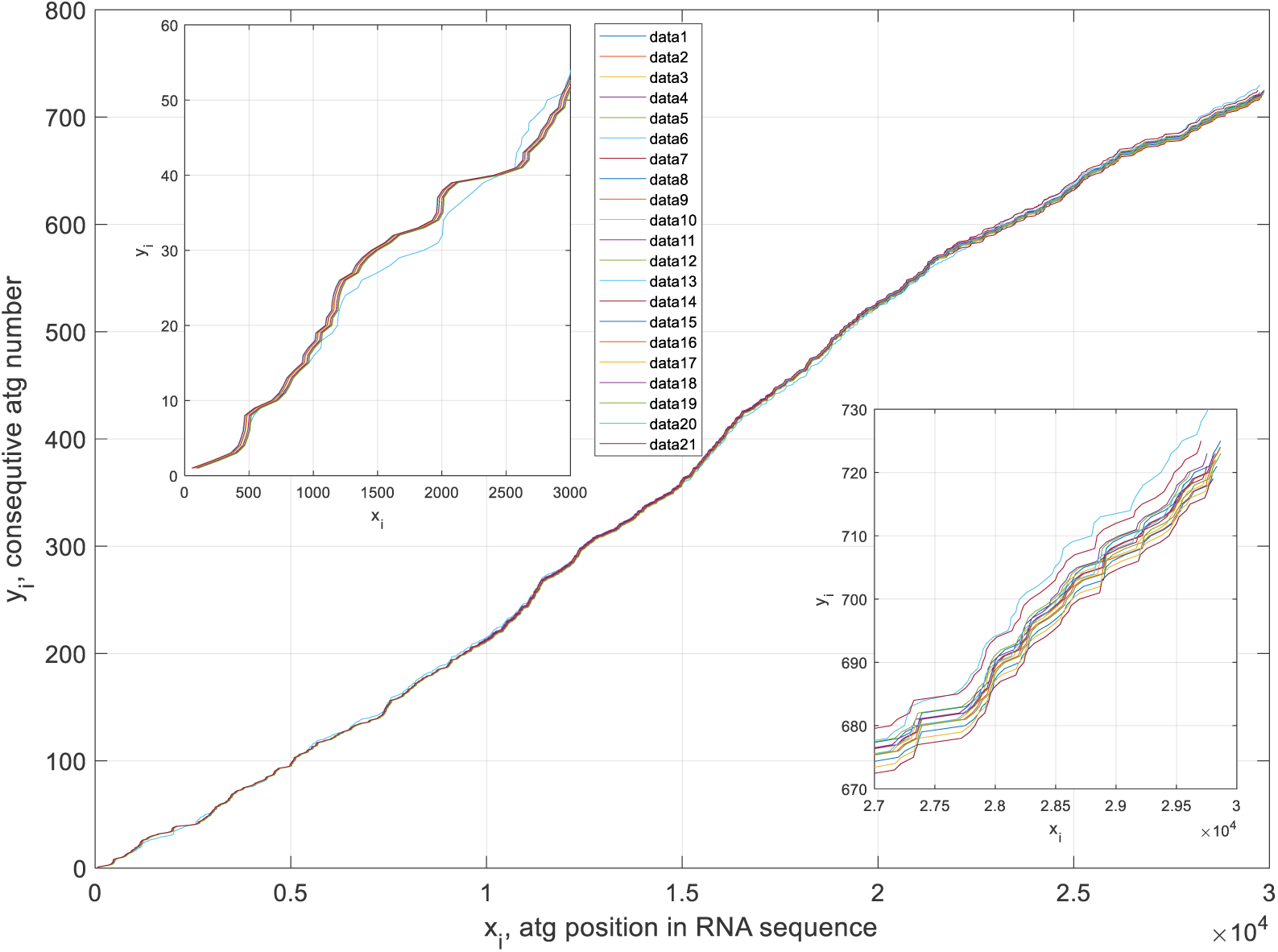
Distributions of *atg*-triplets of 21 SARS Cov-2 complete RNA sequences (rows 1-21, Table 1, Appendix 1). The inlets show these *atg*-distributions at the beginning and end of genome sequences. The numbers of virus *atg*-curves correspond to Table 1, Appendix 1.

Two inlets show the beginning and the tails of these curves to illustrate details. Although, in general, these trajectories are woven firmly, the tails are between the bat’s SARS-CoV-2 light-blue curve (*hCoV-19/bat/Cambodia/RShSTT182/2010*, row 6, Table 1, Appendix 1) and the black trajectory obtained for a sequence from Brazil (*hCoV-19/Brazil/RS-00674HM_LMM52649/2020*, row 14, Table 1, Appendix 1).

A detailed study of each virus from Table 1, Appendix 1 shows that each considered sequence has an individual *atg*-distribution. It means that most mutations are combined with the joint variations of word content, word length and the number of these words. Other mutations with only word content variation may exist. However, the *atg*-walks cannot see them, and the single-symbol distributions considered below will help us to detect these modifications of viruses (See Section 2.2.2.2).

Fig. 6 shows a detailed comparison of samples of five viruses causing increased trouble for specialists with the one from Wuhan, China. The tails of three curves are closed between the Wuhan and Brazil trajectories. The inlets show the details of these curves in their beginning and their end. Although the difference between these curves is not significant, the mutations may have complicated consequences in the rate of contagiousness of viruses.

**Fig. 6.**
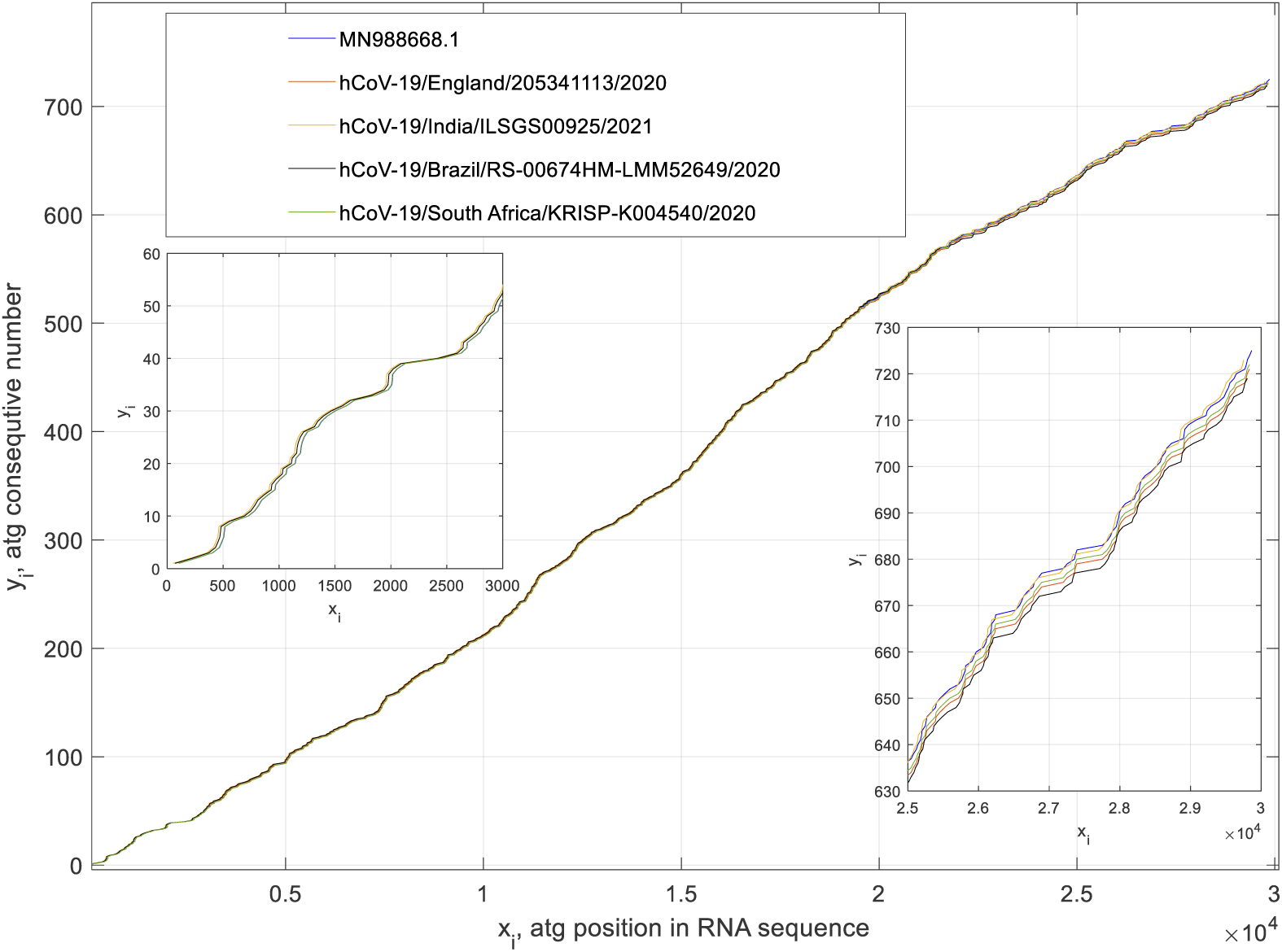
Detailed distributions of *atg*-triplets for five trouble-making SARS Cov-2 complete RNA sequences (rows 1,8,3,14,19, Table 1, Appendix 1). The inlets show these *atg*-distributions at the beginning and end of genome sequences.

There are different techniques for numerical comparing sequences known from data analytics, including, for instance, calculation of correlation coefficients of unstructured data sequences, data distance values, and clustering of data, among others [10],[69]. Researching RNA sequences, we suppose that the error of nucleotide detection is essentially less than one percent; otherwise, the results of comparisons would be instrumentally noisy.

We use a simplified algorithm for quantitative comparing *atg*-distributions of different virus samples. Each numbered *atg*-triplet (*y_i_*) has its coordinate along a sequence (*x_i_*). Thanks to mutations, the length of some coding words varied together with the coordinate (*x_i_*) of a triplet. In our case, we calculated the difference (deviation) between coordinates (*x_i_*) of *atg*-triplets of the same numbers (*y_i_*) in the compared sequences. This operation was fulfilled only for the sequences of the equal number of *atg*-triplets; otherwise, excessive coding words are neglected in comparisons. Of course, such a technique on comparison of geometrical data has its disadvantages. Therefore, if a compared sequence has several *atg*-triplets fewer than the number of *atg*-ones in a reference sequence, the *atg*s of reference RNA are excluded from comparisons. Still, it allows for obtaining some information on mutations of viruses in a straightforward and resultative way that will be seen below.

Our approach supposes choosing a reference nucleotide sequence to compare the genomic virus data of other samples, and it is a complete genomic sequence MN988668.1 from GenBank (row 1, Table 1). Several virus samples from GenBank and GISAID have been studied in this way [42], and some results of comparisons are given in Fig. 7.

**Fig. 7.**
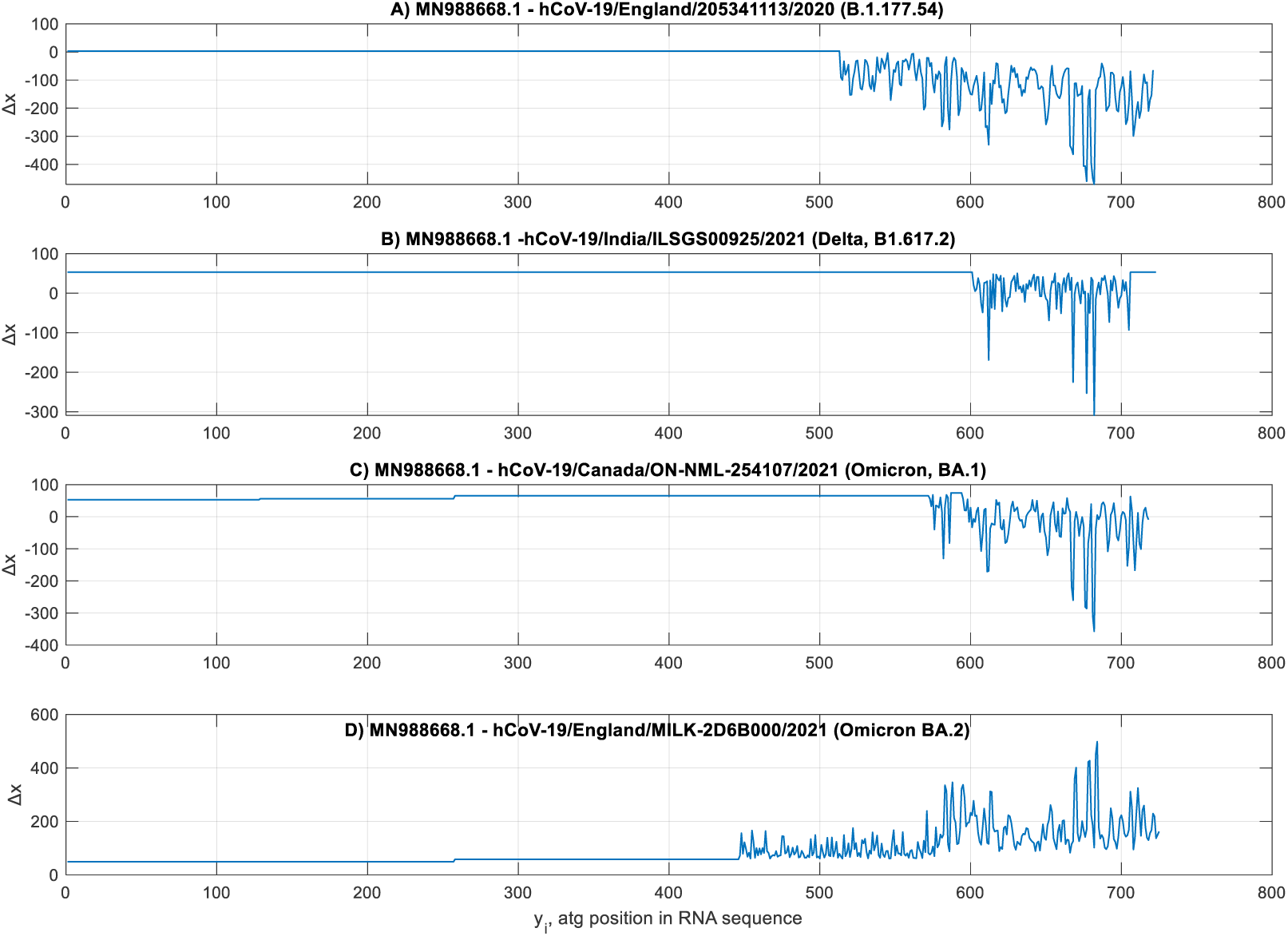
Deviation of *atg*-coordinates in RNAs of four SARS CoV-2 viruses relative to the reference RNA MN988668.1.

The ordinate axis Δ*x* in these plots shows the deviation of coordinates *x_i_* of *atg*-triplets from the *atg*-coordinates of the reference sequence. As a rule, due to the different number of noncoding nucleotides at the beginning of complete RNA sequences, the curves in Fig. 7 have constant biasing along the Δ*x* axis.

The straight parts of these curves mean that the *atg* positions of a compared sequence are not perturbed regarding the corresponding coordinates in the reference RNA. This means that there are no mutations, or they are only with the variation of coding words without affecting their lengths, if these mutations have a place.

In some studied samples (here, and in Refs. [42],[68]), perturbations are near the end of the *orf1ab* gene, as is seen using a graphical tool of GenBank [39]. Perturbations were detected by calculating the *x_i_* coordinates according to known *y_i_*. The *atg*-perturbations could generally occur in any RNA part, considering the random nature of mutations (Fig. 7C, D). Relative deviation |Δ*x_i_*|/*N* did not exceed 1–2% for compared viruses. Although this deviation is mathematically tiny, it may lead to severe consequences in a biological sense.

Our study shows that these difference curves (Fig. 7) are individual for the studied samples. Although mutations without affecting the *atg*-distributions are possible, this individuality, theoretically, may be lost.

There are repeating motifs of comparison curves (Fig. 7 and Refs. [42],[68]). The origin of this is unknown, but it was not coupled with the lineages of viruses and their clades. Other viruses can be studied similarly.

### 3.2. Study of *atg*-Walks of Complete Genome Sequences of the Middle East Respiratory Syndrome-related Coronavirus

The Middle East Respiratory Syndrome-related (MERS) is a viral respiratory illness. The virus’ origin is unknown, but it initially spread through camels and was first registered in Saudi Arabia [70]. Most people infected with the MERS CoV virus developed a severe respiratory disease, which resulted in multiple human deaths.

Our simulation of *atg*-distributions of this virus shows compactness of the calculated curves (Fig. 8, and Table 2, Appendix 1), like the SARS CoV-2 characteristics. It follows that both viruses demonstrate relatively stable features towards the strong mutations connected with the recombination of the virus’s parts. For instance, the divergence of these curves is estimated at around 1% only. On average, the MERS RNAs have fewer *atg*-triplets and longer nucleotide words than the SARS CoV-2 studied sequences.

**Fig. 8.**
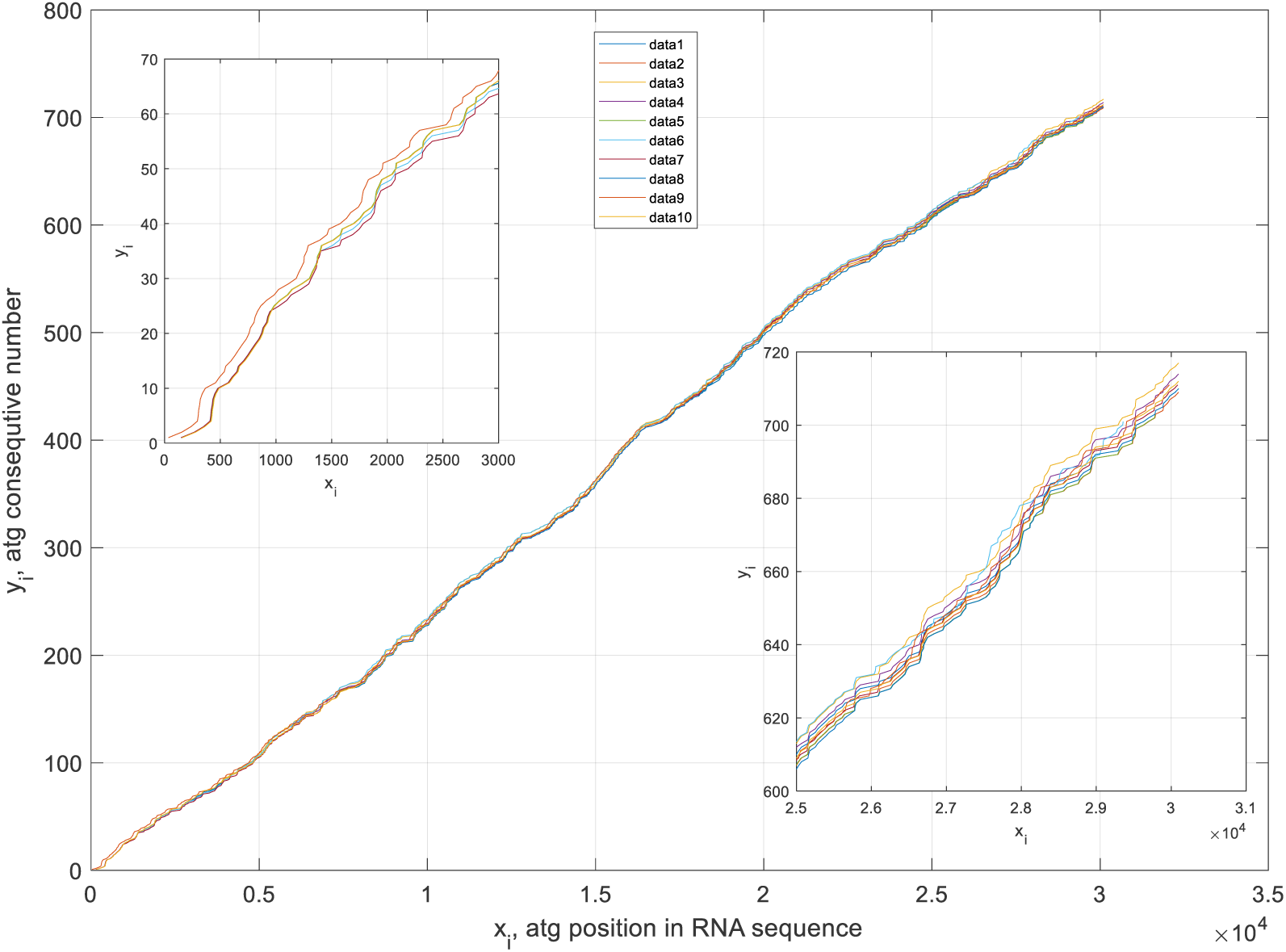
Distributions of *atg*-triplets of ten samples of the MERS CoV complete RNA sequences. Inlet shows the *atg*-distributions at the end of genome sequences (rows 1-10, Table 2, Appendix 1). The inlets offer these *atg*-distributions at the beginning and end of genome sequences. The numbers of virus *atg*-curves correspond to Table 2, Appendix 1.

In general, the two studied coronaviruses (MERS CoV and SARS CoV-2) demonstrate relatively strong stability of their *atg*-distributions towards severe mutations, leading to the variation of codon positions, word lengths and word numbers. This follows the conclusions of many scientists working in virology and virus genomics [71].

### 3.3. Dengue Virus Study

The Dengue virus is spread through mosquito bites. For instance, a recent comprehensive review on the genomics of this virus can be found in Refs. [72],[73]. Unlike the coronaviruses, the Dengue virus (Table 3, Appendix 1) tends to form separate families, i.e. it is less stable compared to SARS CoV-2 and MERS viruses. It has five genotypes (DENV 1–5) and around 47 strains. Only some of them have been studied (below), for which the complete genome data are available from GenBank.

Fig. 9A (rows 1-5, Table 3, Appendix 1) shows the *atg*-distributions of five sequences of Dengue virus-1 found in China. A rather large dispersion of these sequences is seen from these graphs.

**Fig. 9.**
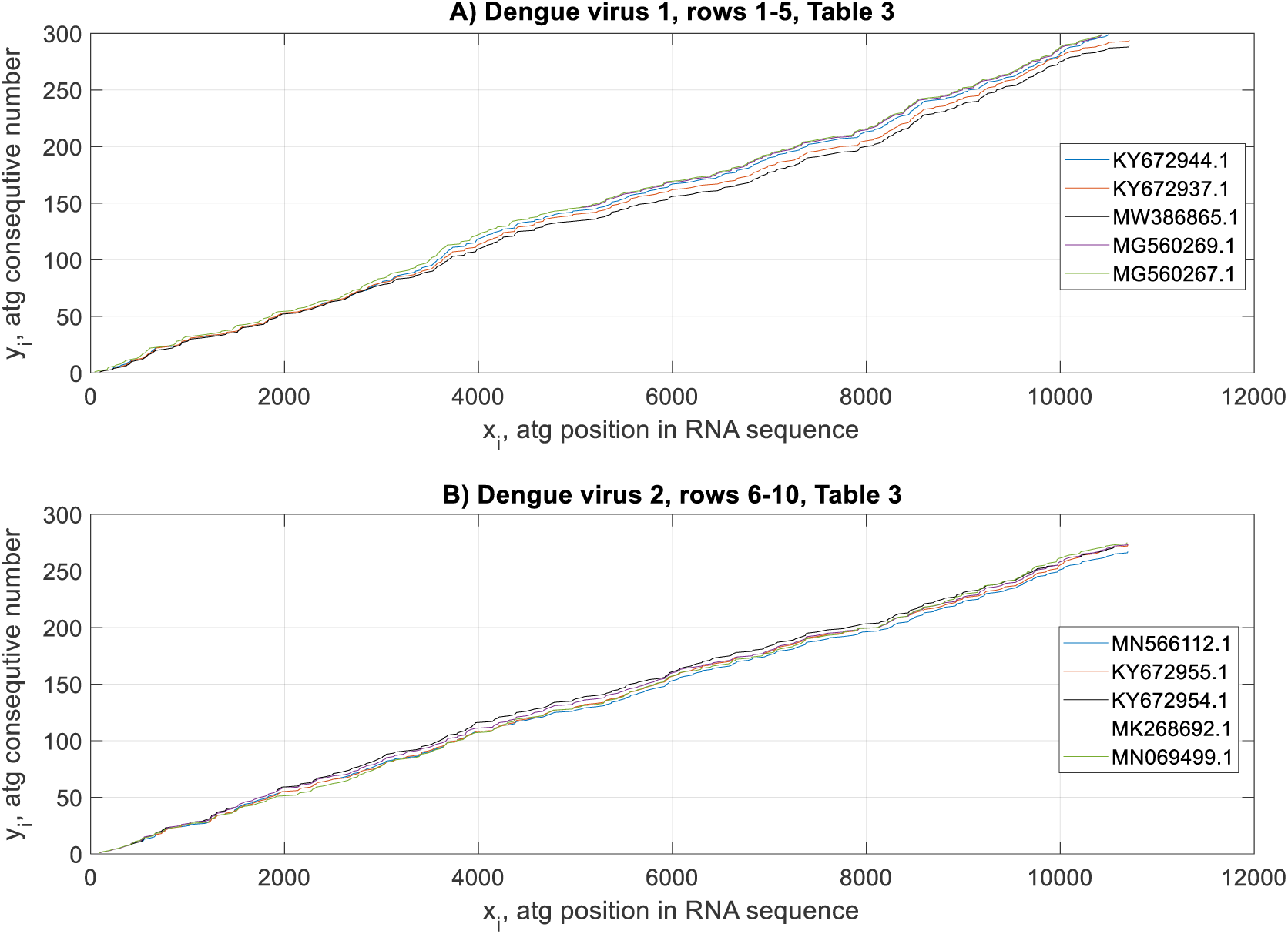
Distributions of *atg*-triplets of complete RNA sequences of the Dengue virus-1 - (A) and Dengue virus-2 - (B). See rows 1-5 and 6-10, correspondingly from Table 3, Appendix 1.

Fig. 9B (rows 6-10, Table 3, Appendix 1) gives the *atg*-distributions of five complete sequences of the Dengue virus-2. These distributions are more compactly localised, although their origin is from different parts of the world. In general, the observed Dengue virus-2 samples have an increased number of shorter words compared to the sequences of Dengue virus-1 (Table 3, Appendix 1).

In Fig. 10A (rows 11-15, Table 3, Appendix 1), five data sets for different strains of Dengue virus-3 registered in many countries are shown. They have about the same number of nucleotides and comparable averaged lengths of words.

**Fig. 10.**
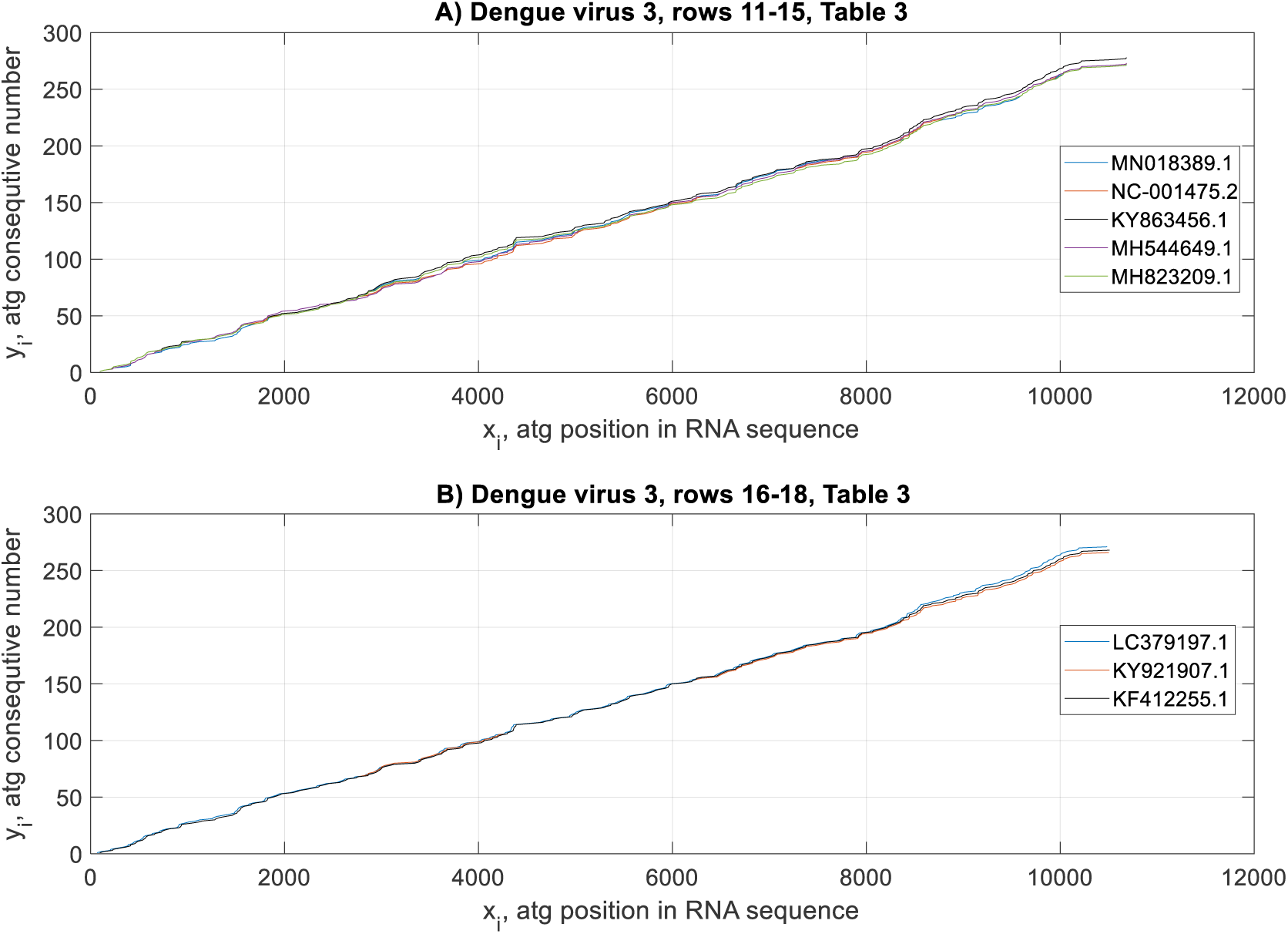
Distributions of *atg*-triplets of complete RNA sequences of Dengue virus-3 (rows 11-15 –(A) and 16-18 –(B) Table 3, Appendix 1).

In Fig. 10B (rows 16-18, Table 3, Appendix 1), three *atg*-distributions of a Gabon-strain [73] of Dengue virus-3 are given. It is supposed that this strain mutated from the earlier registered Gabon Dengue virus lines (Fig. 10A). However, they are different in the length of complete genome sequences and their statistical characteristics, which are considered in Section 3.5 below.

Fig. 11A (rows 19-20, Table 3, Appendix 1) shows the *atg*-distributions of two Gabon-originated Dengue viruses that can relate to predecessors of other Dengue viruses of this family. Fig. 11B (rows 21-25, Table 3, Appendix 1) presents the *atg*-distributions of complete RNA sequences for five Dengue virus-4 samples. They have individual and statistical differences with the above-considered Dengue viruses.

**Fig. 11.**
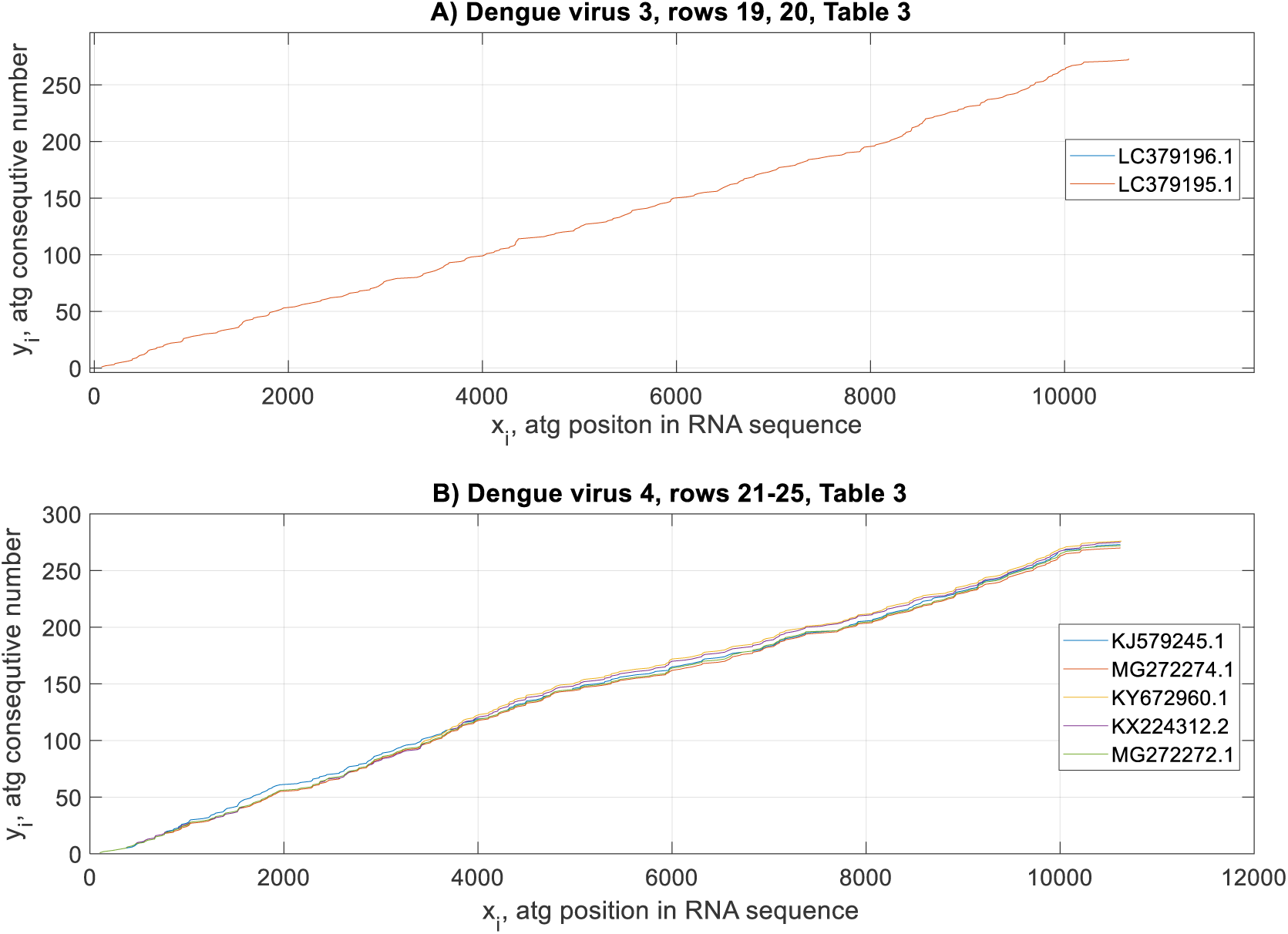
Distributions of *atg*-triplets of complete RNA sequences of Dengue virus-3 (rows 19, 20 - (A), Table 3, Appendix 1) and Dengue-4 (rows 21-25 - (B), Table 4, Appendix 1).

A consolidated plot of all *atg*-curves of the Dengue RNAs studied here is shown in Fig. 12. There is substantial divergence of these trajectories in agreement with the mutation rate of this virus being relatively strong. For instance, this deviation estimated at *x_i_* = 10000 is 14.1%.

**Fig. 12.**
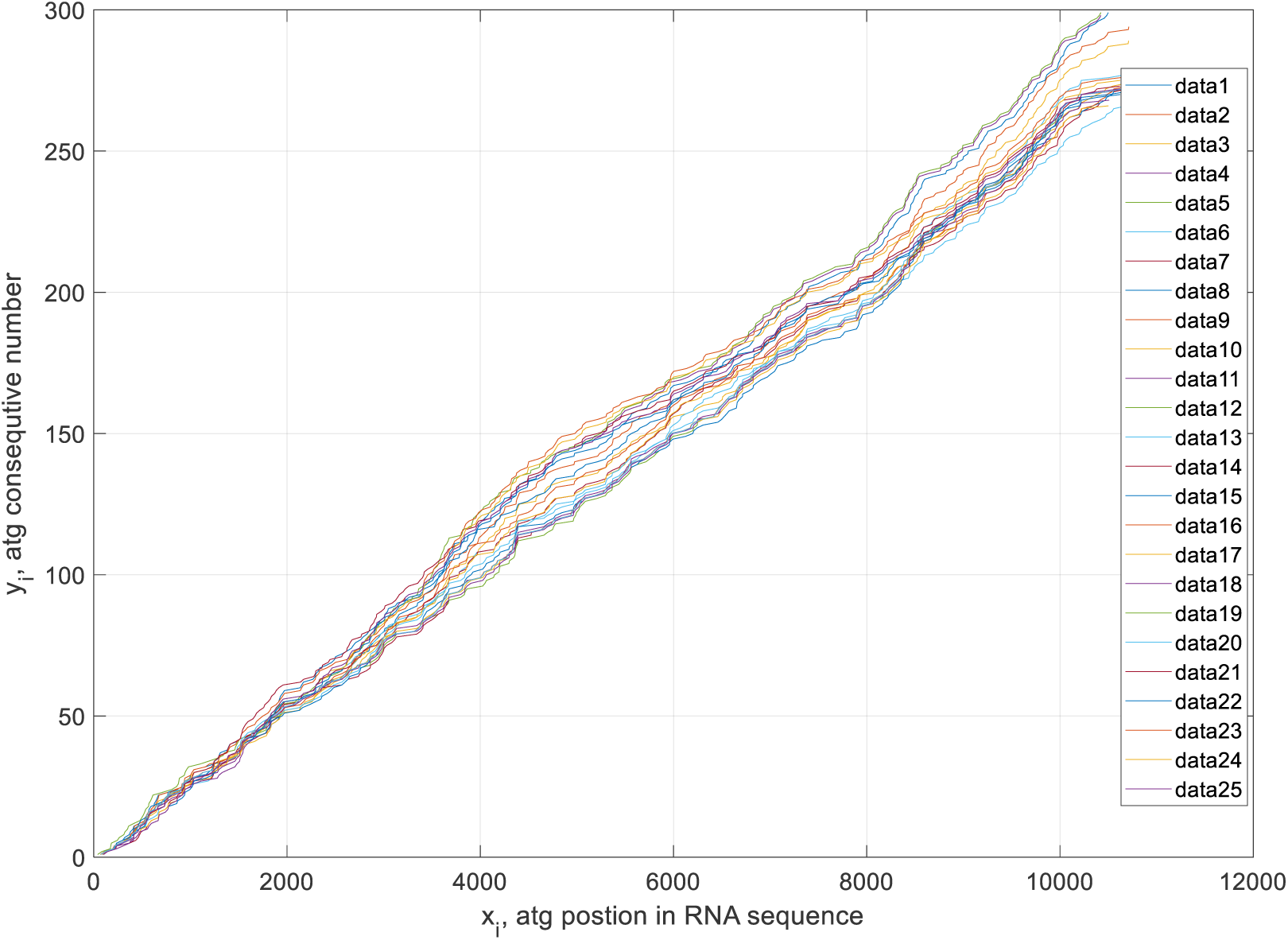
Consolidated picture for all Dengue virus samples studied in this paper. The numbers of virus *atg*-curves correspond to Table 3, Appendix 1.

### 3.4. Analysis of *atg*-Walks of Complete Genome Sequences of the Ebola Virus

There are four strains of Ebola virus known in the world, although many other mutations of this virus can be found. Like the Dengue virus, the Ebola virus shows instability and an increased rate of mutations. Initially, the infection was registered in South Sudan and the Democratic Republic of the Congo, and it spreads due to contact with the body fluids of primates and humans. This fever is distinguished with a high death rate (from 25% to 90% of the infected individuals). A recent comprehensive review on the genomics of this virus can be found in Ref. [74].

The Ebola virus RNA consists of 19 000 nucleotides and more than three hundred *atg*-triplets. Fig. 13A shows four sequences of this virus belonging to the EBOV strain registered from Zaire and Gabon. Three of them are very close to each other, but the mutant Zaire virus (in red) has some differences from the three others. The samples collected in Sudan (SUDV) are closer to each other (Fig. 13B), but they have an increased number of *atg*-triplets and shorter words.

**Fig. 13.**
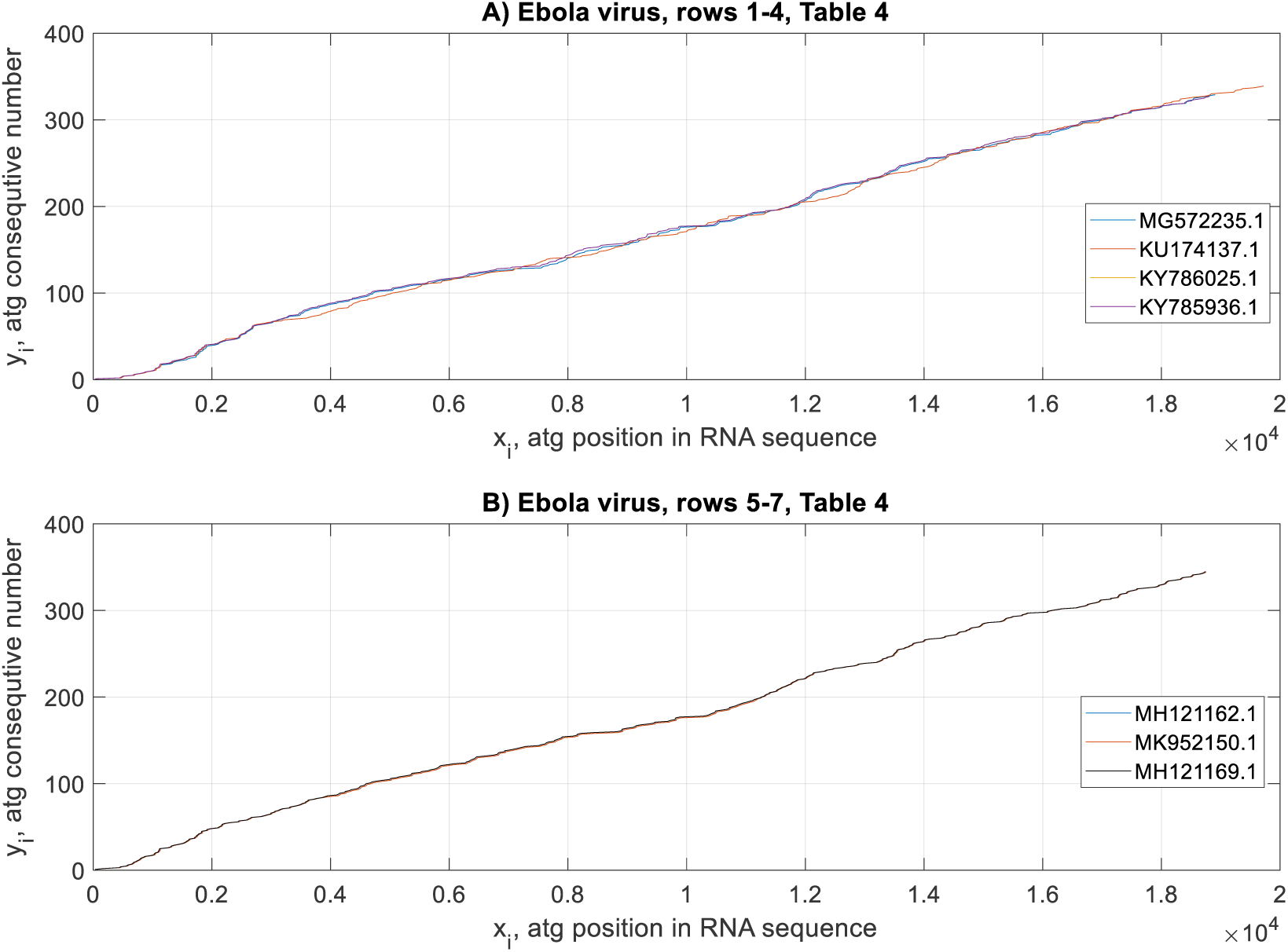
Distributions of *atg*-triplets of complete RNA sequences of the Ebola - EBOV) virus from Zaire and Gabon (rows 1-4 - (A), Table 4, Appendix 1) and Ebola virus - SUDV from Sudan (rows 5-7 - (B), Table 4, Appendix 1).

The Bombali virus is considered a new strain of the Ebola virus registered in Sierra Leone, West Africa. The *atg*-distributions of the five RNA sequences studied here are different even visually from the two reviewed above, as seen in Fig. 14A. Another Ebola virus strain that can be compared with the one studied above is the Bundibugyo (BDBV) virus, whose three *atg*-distributions are shown in Fig. 14B.

**Fig. 14.**
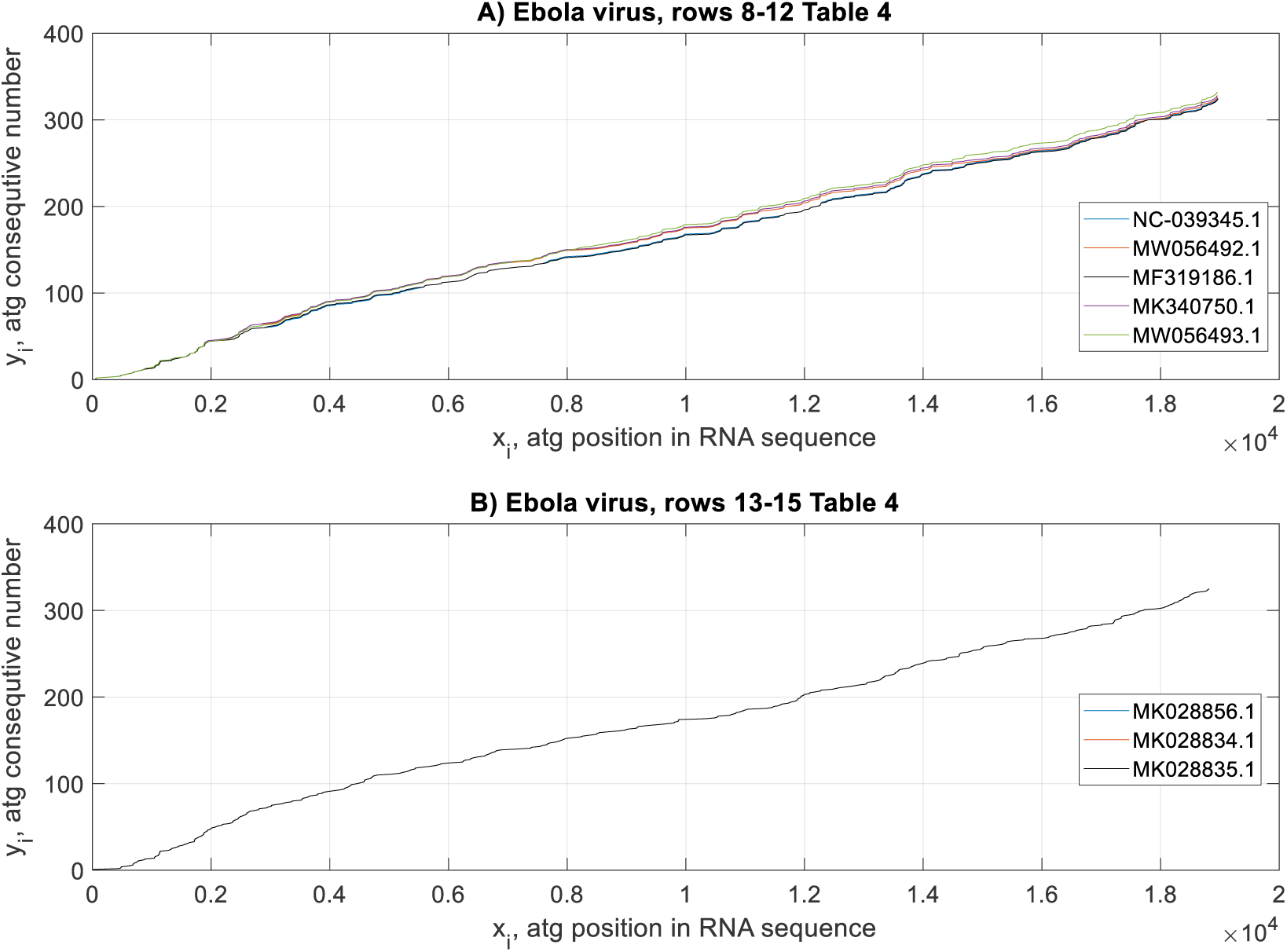
Distributions of *atg*-triplets of five complete RNA sequences of the Ebola (Bombali) virus (rows 8-12 – (A), Table 4, Appendix 1) and three complete RNA sequences of the Ebola Bundibugyo (BDBV) virus. See rows 13-15 – (B), Table 4, Appendix 1.

The calculated distributions are consolidated in Fig. 15 to compare all four strains, where, instead of points, the results are represented by thin curves to make these distributions more visible. Here, the tendency of *atg*-curves to divergence and forming clusters is seen. For instance, the deviation of strains estimated at *x_i_* = 18000 is around 9%.

**Fig. 15.**
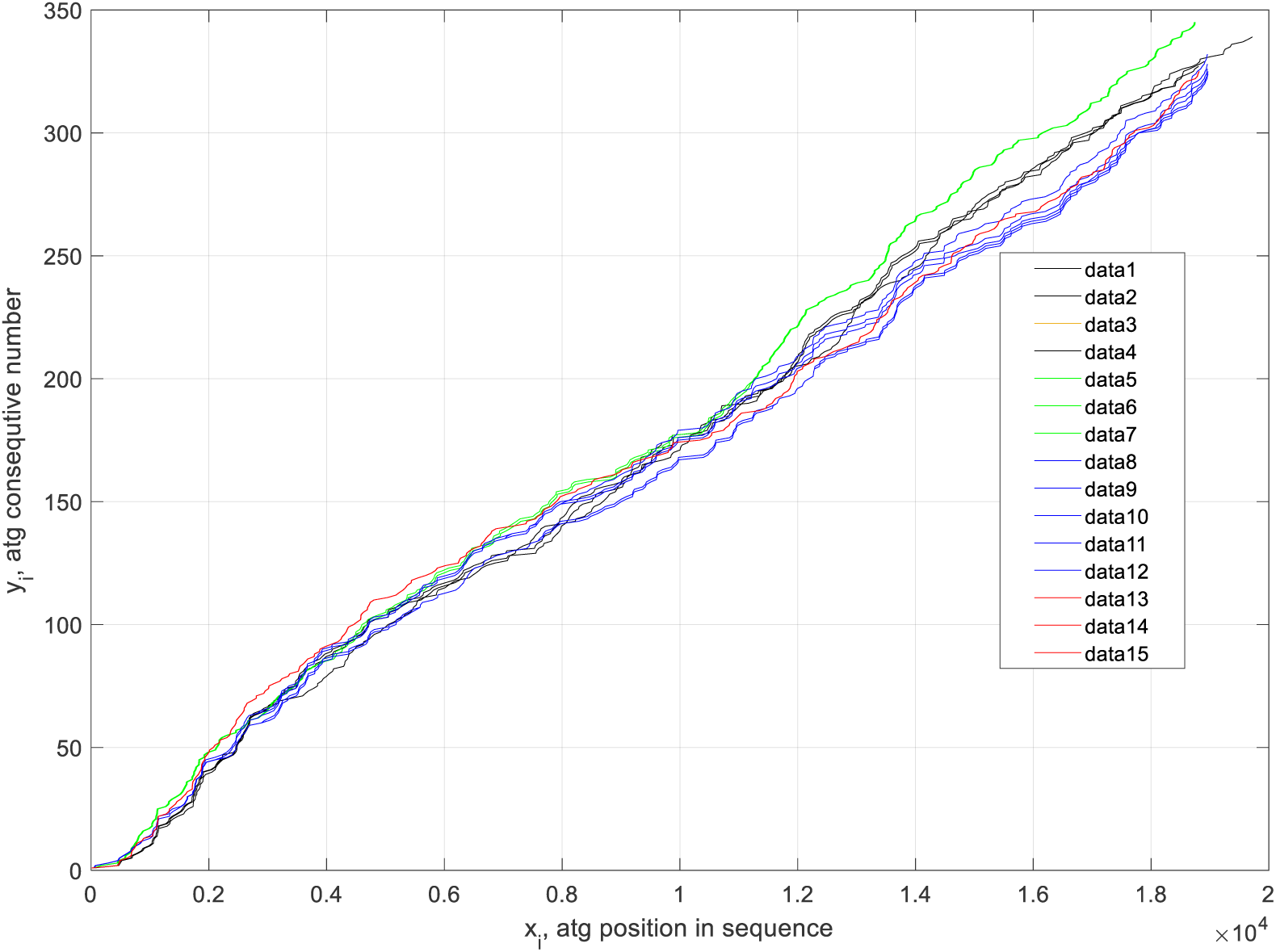
Consolidated representation of *atg*-distributions of four strains of the Ebola virus. Black colour – EBOV; Green colour – SUDV; Violet colour – Bombali; Red colour – BDBV. The numbers of virus *atg*-curves correspond to Table 4, Appendix 1.

Reviewing all above-obtained results, the *atg*-walk is an effective visualisation tool sensitive to the viral RNA mutations connected with the number of codons’ variation, word width, and *atg*-coordinates. It allows to detect the viruses with essentially unstable genomes distinguished by their increased deviation of *atg*-walks and their fractal properties.

### 3.5. Statistical Characterisation of *atg*-Walks: Calculating, Mapping, and Processing of the Inter-*atg* Distance Values

In this research, after applying the above-mentioned tool FracLab (See Section 2.2.3), it was discovered that all studied genomic sequences of the SARS CoV-2, MERS CoV, Dengue and Ebola viruses have fractality in their word-length distributions. The fractal regularisation dimension values [60, 61] calculated by us correlated with uniform linear ideal polymers [25]. The results on this matter are placed in columns 6 of Tables 1-4, Appendix 1.

Fig. 16 shows the fractal dimension values of 20 genome sequences of SARS CoV-2 viruses in humans and one found in bats (Table 1, Appendix 1). Ten samples of MERS CoV genome sequences (Table 2, Appendix 1) are given in the same figure. Although the *atg*-distributions of these viruses are visually close to each other, the word-length fractal dimension values were essentially different.

**Fig. 16.**
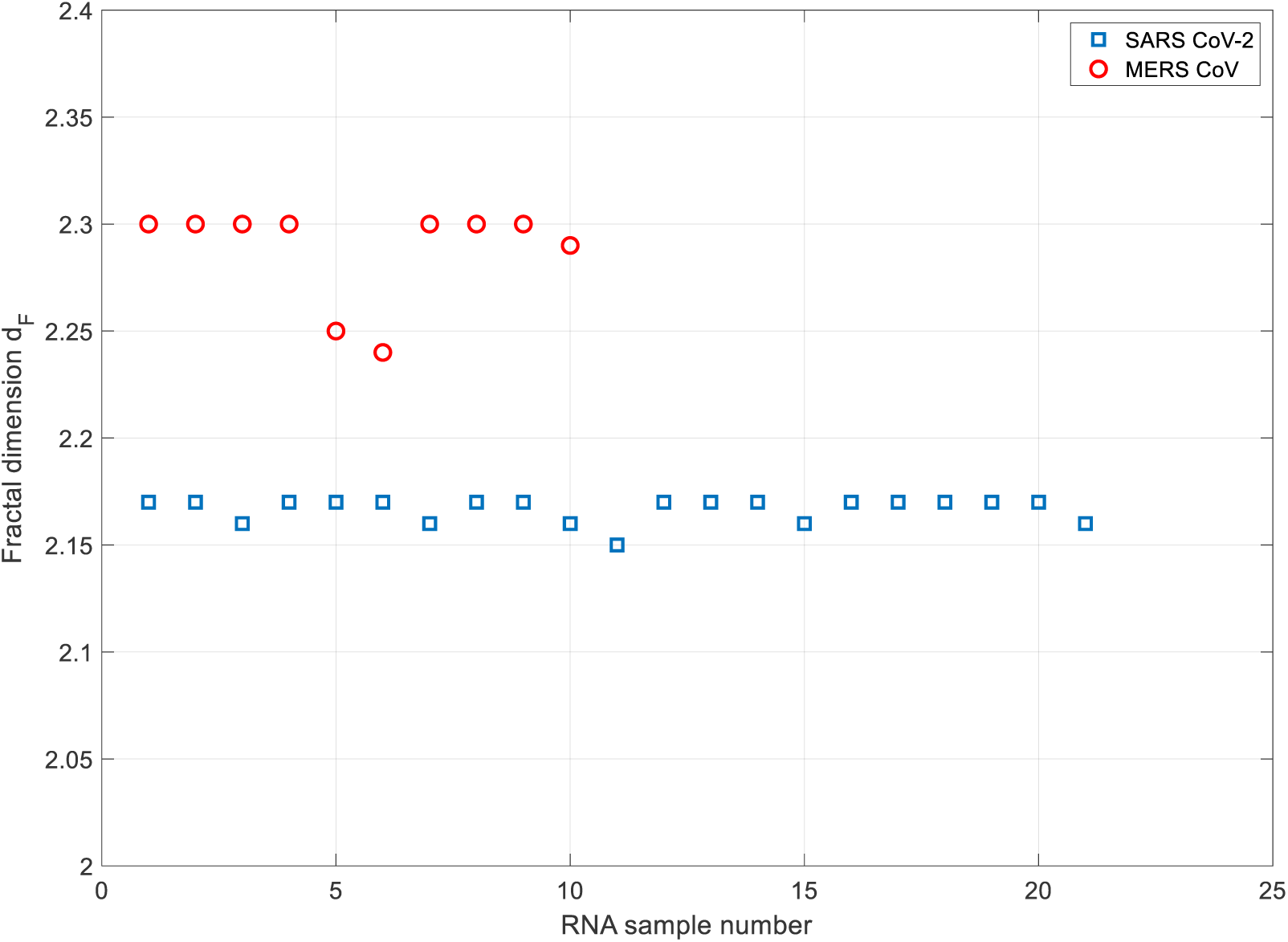
Fractal dimensions *d_F_* of word-length 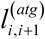 distributions of complete genome sequences of the SARS CoV-2 and MERS CoV viruses. The number of samples corresponds to Tables 1 and 2 of Appendix 1.

The Dengue virus has five families and 47 strains; they have different *atg*-distributions and fractal dimensions. Some strains are close to each other according to the fractal calculations (Fig. 17). This gives a reason to conclude that the RNAs of the considered strains have similarities in the *atg*-distributions.

**Fig. 17.**
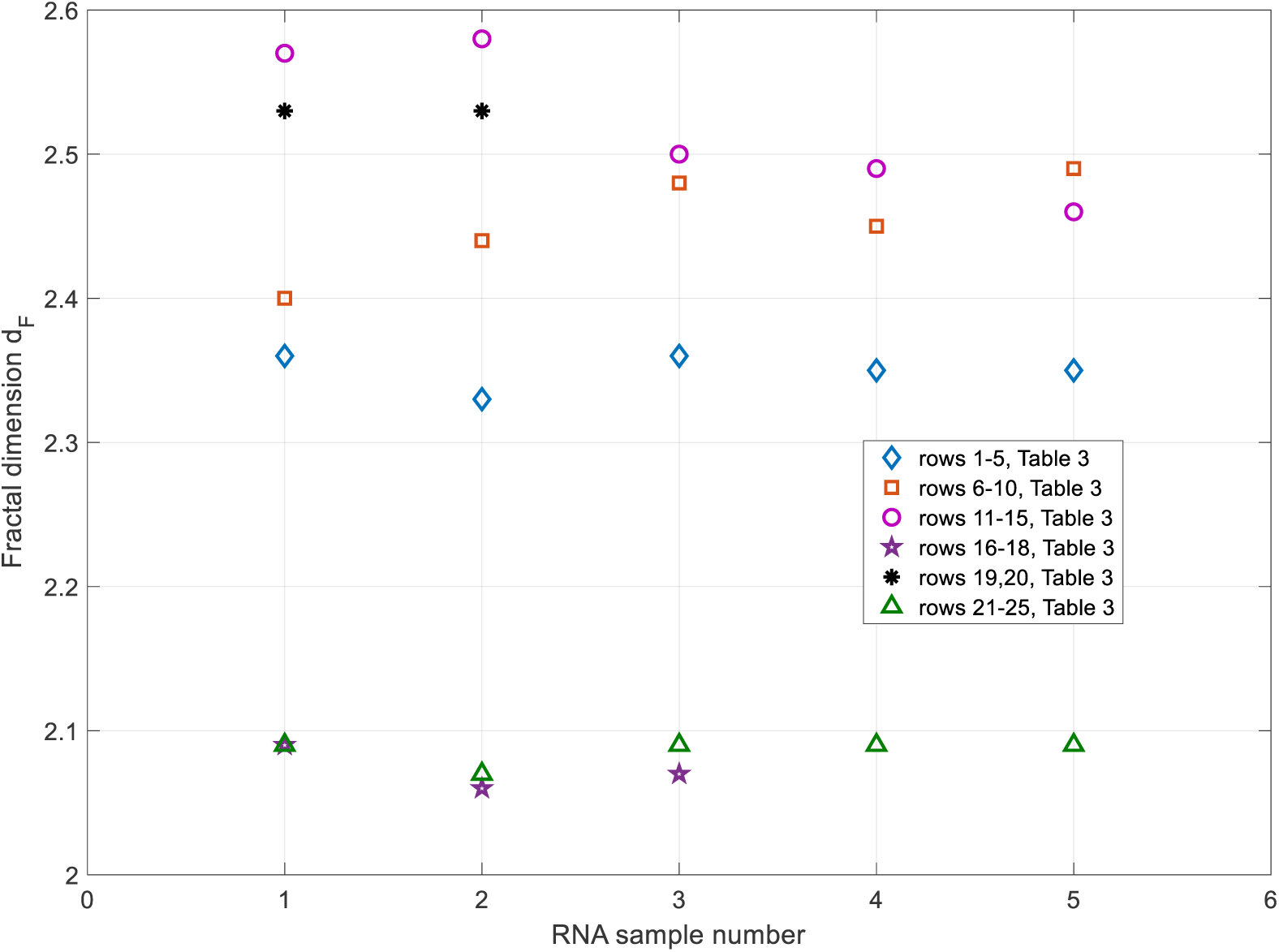
Fractal dimensions *d_F_* of word-length 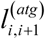 distributions of complete genome sequences of the Dengue 1-4 viruses and their strains.

The same conclusion is evident in Fig. 18, where the fractal dimensions of several strains of the Ebola virus are given.

**Fig. 18.**
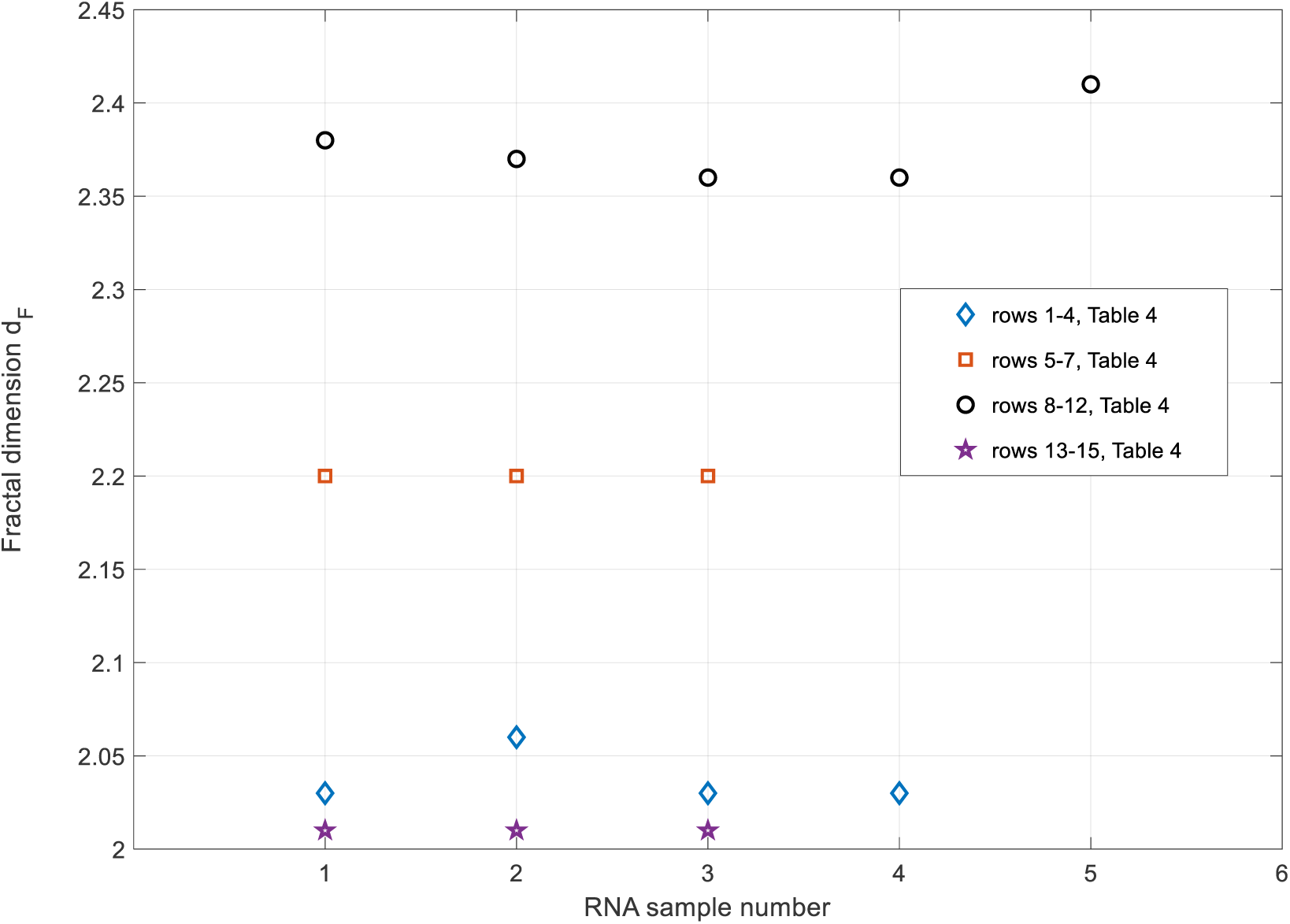
Fractal dimensions *d_F_* of word-length 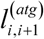 distributions of complete genome sequences of the Ebola virus strains.

An analysis of the statistical characteristics of *atg*-distributions shows that the average word length is coupled in a certain way with the fractal dimension. As a rule, the word-length reduction increases the fractal dimension, which means a more complicated distribution of *atg*-triplets.

A comparative analysis of Fig. 16 gives us that the RNAs of some viruses of the same strain have a somewhat stable fractal dimension value *d _F_*. This is typical for the studied samples of the SARS CoV-2 and MERS CoV viruses.

It is known that the Dengue and Ebola viruses have increased mutation rates. If we follow the contemporary classification of these viruses, they have essentially different fractal dimension values even for the samples belonging to the same strain (Fig. 17, Dengue 3 and Fig. 18, Ebola), which points to the increased variability of these viruses.

Concluding the research on fractal properties of the studied viruses, it is worthwhile to say that fractal dimension is coupled with the complexity of a polymer design, and this complexity can define the rate of chemical reaction with these polymers. Besides, a limited value of this dimension shows on spatial correlation in nucleotide distribution. In our cases, additionally, sample-to-sample large variation of the dimension is with unstability of virus species. See, for instance, MERS/SARS CoV-2 and Dengue/Ebola viruses.

## 4. Discussion

The research on the RNAs and DNAs of viruses and cellular organisms is a highly complex problem because of the many nucleotides of these organic polymers, unclear mechanisms of their synthesis and pathological mutation consequences for host organisms. Although many mathematical tools have been developed, new studies are exciting and can be fruitful.

In this paper, the viral RNAs were studied using a novel algorithm based on exploring RNA patterns of arbitrary length. One of the operations of this algorithm is with the numerical mapping of RNA characters, which is performed by calculating the Hamming distance between the preliminary binary-expressed queries and RNA symbols. This allows fulfilling these steps approximately twice as fast regarding the operations with real numbers [43]. The results of the application of this algorithm are verified by comparing them with complete RNA sequences. Considering that this algorithm can search arbitrary-length patterns, the trajectories of separate symbols can be combined with the *atg*-walks for multi-scale plotting and analysis of RNAs, as shown in this contribution.

The mentioned codon-starting *atg*-triplets compose relatively stable 1-D distributions called the RNA schemes in this paper. These distributions have been studied using our algorithm applied to complete RNA sequences of the SARS CoV-2, MERS CoV, Dengue and Ebola viruses registered in GenBank [39] and GISAID [40].

The following properties of virus RNAs have been found in our research and not seen earlier:

1. The relative stability of *atg*-schemes towards intra-family mutations when the geometry of *atg*-curves is only slightly distorted (Section 3)
2. The highly compact *atg*-curves of the SARS CoV-2 and MERS CoV viruses despite their continuous mutation (to the date of submission), estimated visually and quantitively (Sections 3.1 and 3.2)
3. More substantial divergence of *atg*-curves of the Dengue and Ebola viruses in comparison to SARS CoV-2 and MERS CoV (Sections 3.1, 3.2, 3.3 and 3.4)
4. Tendency towards clustering *atg*-curves in the limits of one virus family (Ebola case, Section 3.4)
5. Distribution of single RNA’s symbols and *atg*-triplets according to the random fractal Cantor rule (Section 2.2.2.1)
6. Possible global correlation of the inter-triplet distances due to this fractality (Section 2.2.3)
7. Correlation of dispersion of fractal dimension values of *atg*-distributions of genomes with the instability of viruses (Section 3.5)

## 5. Conclusions

In this paper, the visual and quantitative analyses of viral RNAs were performed using a novel algorithm to calculate the RNA pattern positions in the studied sequences. A part of this code uses binary symbols of RNA nucleotides for accelerated search. The algorithm allows more effective genomic studies by building 1-D distributions of different patterns and combining these sets on a single multi-scale plot.

The proposed techniques were applied to analyse SARS CoV-2 and MERS CoV as well as the Dengue and Ebola viruses. The 1-D distributions of codon starting *atg*-triplets (*atg*-schemes of RNAs) were calculated and plotted for these species. The levels of stability of these distributions were estimated numerically, including calculation of fractal dimensions of these sets and analysis of these results. Particularly, deviations of *atg*-distributions and their statistical characteristics (fractal dimension values) show on stronger stability of the first viruses in comparison to the Dengue and Ebola species that makes more hopes on possible reliable control of corona viruses spread in the future.

The developed approach is interesting in the study of mutation of viruses and building their phylogenetic trees. Further developments are needed in applying this approach to study large genomic sequences and proteins, including code accelerating and minimisation of required memory. The applicability of the algorithm will be increased, enhancing this code by an interactive means showing the affiliation of different nucleotides and *atg*-triplets to specific genes.

## Abbreviations

RNA: Ribonucleic acid
DNA: Deoxyribonucleic acid
SARS CoV-2: Severe Acute Respiratory Syndrome Coronavirus 2
cDNA: complementary DNA
GISAID: Global Initiative on Sharing All Influenza Data
NP: nondeterministic polynomial
UTF-8: Unicode Transformation Format-8 bit
*US-ASCII*: American Standard Code for Information Interchange
MERS CoV: Middle-East Respiratory Syndrome-related Corona Virus.

## Acknowledgments

The authors thank the GenBank® [39] and GISAID [40] genetic data banks, and all researchers placed their genomic sequences in them. The online text processing service of https://onlinetexttools.com/ is appreciated.

## Authors’ contributions

All authors are contributed equally

## Declarations

### Ethical approval and consent to participate

Not applicable

### Consent for publication

Not applicable

## Appendix 1: Results of statistical characterisation of complete genetic sequences of the SARS CoV-2, MERS CoV, Dengue and Ebola viruses

**Table 1.** Severe acute respiratory syndrome coronavirus 2, (GenBank, GISAID), *atg*-walk

| # | GenBank or GISAID Virus Name, Clade, Lineage, Registration Year, Sequencing Technology | Number of Nucleotides in the Sequence | Number of *atg*-Triplets in the sequence | Word Median Length | RMS Word Length | Fractal Dimension of the Word-length Distribution |
| --- | --- | --- | --- | --- | --- | --- |
|  | 1 | 2 | 3 | 4 | 5 | 6 |
| 1 | GenBank: *MN988668.1*, Severe acute respiratory syndrome coronavirus 2 isolate 2019-nCoV WHU01, Wuhan, China, 2020, Illumina | 29881 | 725 | 29 | 57.93 | 2.17 |
| 2 | *hCoV-19/Japan/NGY-NNH-075/2021*, GR, B.1.1.64, Illumina MiSeq, Sanger | 29848 | 722 | 29 | 58.03 | 2.17 |
| 3 | *hCoV-19/India/ILSGS00925/2021*, G, (Delta) B.1.617.2, Illumina NextSeq550 | 29782 | 723 | 28.05 | 57.77 | 2.16 |
| 4 | *hCoV-19/South Korea/KDCA3504/2021*, GH, B.1.497, Illumina Miseq | 29901 | 722 | 29 | 57.9602 | 2.17 |
| 5 | *hCoV-19/Taiwan/TSGH-34/2020*, S, A.1, Illumina NovaSeq4000 | 29903 | 724 | 29 | 57.79 | 2.17 |
| 6 | *hCoV-19/bat/Cambodia/RShSTT182/2010*, A.1, (bat virus), 2021, Illumina NextSeq | 29787 | 730 | 29 | 55.81 | 2.17 |
| 7 | *hCoV-19/Austria/CeMM3224/2021*, GR, B.1.1.244, Illumina NovaSeq | 29782 | 721 | 30 | 59.03 | 2.16 |
| 8 | *hCoV-19/England/205341113/2020*, GV, B.1.177.54, Illumina NextSeq | 29862 | 721 | 29 | 57.97 | 2.17 |
| 9 | *hCoV-19/Ireland/D-NVRL-e84IRL94434/2021*, GV, B.1.177, Illumina | 29523 | 719 | 29 | 59.56 | 2.17 |
| 10 | *hCoV-19/Netherlands/UT-RIVM-13868/2021*, GH, B. 1.160, Nanopore MinION | 29782 | 720 | 28 | 58.17 | 2.16 |
| 11 | *hCoV-19/Norway/0179/2021*, GH, B.1.36, Nanopore Gridlon | 29782 | 723 | 28 | 57.88 | 2.15 |
| 12 | *hCoV-19/Russia/IVA-CRIE-L188N0202/2021*, GR, B.1.1.317, Illumina | 29735 | 720 | 29 | 57.77 | 2.17 |
| 13 | *hCoV-19/Spain/RI-IBV-99016064/2021*, GV, B.1.221, Illumina MiSeq | 29865 | 719 | 29 | 59.56 | 2.17 |
| 14 | *hCoV-19/Brazil/RS-00674HM_LMM52649/2020*, GR, B.1.1.33, Illumina Miseq | 29867 | 719 | 29 | 58.31 | 2.17 |
| 15 | *hCoV-19/Canada/ON-S2383/2021*, GH, B. 1.36.38, Illumina MiniSeq | 29830 | 722 | 29 | 57.89 | 2.16 |
| 16 | *hCoV-19/Mexico/CMX-INER-0222/2020*, G, B.1.551, Illumina NextSeq | 29885 | 724 | 29 | 57.83 | 2.17 |
| 17 | *hCoV-19/USA/TX-HHD-2102044112/2021*, GR, B.1.1.244, Illumina MiSeq | 29819 | 720 | 29 | 58.10 | 2.17 |
| 18 | *hCoV-19/USA/CA-LACPHL-AF00513/2021*, GH, B.1.429, Illumina MiSeq | 29844 | 723 | 29 | 57.86 | 2.17 |
| 19 | *hCoV-19/South Africa/KRISP-K004540/2020*, GR, B.1.1.56, Illumina MiSeq | 29851 | 722 | 29 | 57.90 | 2.17 |
| 20 | *hCoV-19/Canada/ON-NML-254107/2021*, GR, BA.1 (Omicron), Oxford Nanopore GridION | 29685 | 718 | 29 | 57.73 | 2.17 |
| 21 | *hCoV-19/England/MILK-2D6B000/2021*, GRA, BA.2 (Omicron), Illumina NovaSeq | 29724 | 725 | 28.5 | 57.48 | 2.16 |

**Table 2.** The Middle East respiratory syndrome-related coronavirus, (GenBank), *atg*-walk

| # | GenBank Virus Name and Accession Number, Registration Year, Sequencing Technology | Nucleotides Number | Number of atg-Triplets | Word Median Length | RMS Word Length | Fractal Regularization Dimension of the Word-length Distribution |
| --- | --- | --- | --- | --- | --- | --- |
|  | 1 | 2 | 3 | 4 | 5 | 6 |
| 1 | *MF598617.1*, Middle East respiratory syndrome-related coronavirus strain camel/UAE_B25_2015, United Arabian Emirates, AE, 2017, Illumina; Sanger dideoxy sequencing | 30123 | 712 | 30 | 58.8 | 2.30 |
| 2 | *MF598595.1*, Middle East respiratory syndrome-related coronavirus strain camel/UAE_B2_2015, United Arabian Emirates, 2017, Illumina; Sanger dideoxy | 30123 | 709 | 30 | 59.04 | 2.30 |
| 3 | *NC-019843.3*, Middle East respiratory syndrome-related coronavirus isolate HCoV-EMC/2012, Saudi Arabia, 2020, Sanger dideoxy | 30119 | 717 | 30 | 58.48 | 2.30 |
| 4 | *KY673148.1*, Middle East respiratory syndrome-related coronavirus strain Hu/Oman_50_2015, 2017, Sanger dideoxy | 30123 | 714 | 29 | 58.74 | 2.30 |
| 5 | *KT225476.2*, Middle East respiratory syndrome coronavirus isolate MERS-CoV/THA/CU/17_06_2015, Oman/Thailand, 2017, Sanger dideoxy | 29809 | 703 | 30 | 59.03 | 2.25 |
| 6 | *MG923479.1*, Middle East respiratory syndrome-related coronavirus isolate MERS-CoV camel/Nigeria/NV1712/2016, 2018, Sanger dideoxy | 29455 | 701 | 30 | 58.08 | 2.24 |
| 7 | *MK967708.1*, Middle East respiratory syndrome-related coronavirus isolate Merscov/Egypt/Camel/AHRI-FAO-1/2018, 2019, CLC genomic workbench | 30106 | 711 | 30 | 58.05 | 2.30 |
| 8 | *MT361640.1*, Mutant Middle East respiratory syndrome-related coronavirus clone MERS-CoV YKC, South Korea, 2021, sequencing technology is described in [75] | 30136 | 710 | 30 | 58.90 | 2.30 |
| 9 | *KT326819.1*, Middle East respiratory syndrome coronavirus strain MERS-CoV/KOR/KNIH/001_05_2015, South Korea, 2017, Illumina and Sanger dideoxy | 29995 | 711 | 30 | 58.86 | 2.30 |
| 10 | *MK129253.1*, Middle East respiratory syndrome-related coronavirus isolate MERS-CoV/KOR/KCDC/001_2018-TSVi, South Korea, 2019, Sanger dideoxy | 30150 | 712 | 30 | 58.81 | 2.29 |

**Table 3.** The Dengue virus, (GenBank), *atg*-walk

| # | GenBank Virus Name, Registration Year, Sequencing Technology | Nucleotides Number | Number of *atg*-Triplets | Word Median Length | RMS Word Length | Fractal Regularization Dimension of the Word-length Distribution |
| --- | --- | --- | --- | --- | --- | --- |
|  | 1 | 2 | 3 | 4 | 5 | 6 |
| 1 | KY672944.1, Dengue virus 1 isolate DENV-1/China/YN/YNH22 (2013), 2019, Sanger dideoxy | 10709 | 299 | 23 | 47.74 | 2.36 |
| 2 | KY672937.1, Dengue virus 1 isolate DENV-1/China/YN/DGRL-6(2014), 2019, Sanger dideoxy | 10738 | 294 | 23 | 50.02 | 2.33 |
| 3 | MW386865.1, Dengue virus 1 isolate YNBN04, China, 2020, Sanger dideoxy | 10742 | 289 | 24 | 50.81 | 2.36 |
| 4 | MG560269.1, Dengue virus 1 isolate P1253/China/GD/CZ/2014, 2018, Sanger dideoxy | 10583 | 298 | 23 | 47.55 | 2.35 |
| 5 | MG560267.1, Dengue virus 1 isolate P1258/China/GD/CZ/2014, 2018, Sanger dideoxy | 10583 | 299 | 23 | 47.22 | 2.35 |
| 6 | MN566112.1, Dengue virus 2 isolate New Caledonia-2018-AVS127, 2020, Illumina | 10722 | 267 | 32 | 52.24 | 2.4 |
| 7 | KY672955.1, Dengue virus 2 isolate DENV-2/China/YN/15DGR65(2015), 2019, Sanger dideoxy | 10723 | 273 | 28 | 52.77 | 2.44 |
| 8 | KY672954.1, Dengue virus 2 isolate DENV-2/China/YN/JH1516(2015), 2019, Sanger dideoxy | 10665 | 271 | 29 | 51.50 | 2.48 |
| 9 | MK268692.1, Dengue virus 2 isolate DENV-2/TH/1974, Thailand, 2019, Sanger dideoxy | 10721 | 274 | 28 | 52.67 | 2.45 |
| 10 | MH069499.1, Dengue virus 2 strain DENV-2/VE/IDAMS/910105, Venezuela, 2018, Illumina | 10712 | 275 | 28 | 52.84 | 2.49 |
| 11 | MN018389.1, Dengue virus 3 isolate D17011, China, 2020, Sanger dideoxy sequencing | 10708 | 272 | 28 | 55.46 | 2.57 |
| 12 | NC_001475.3, Dengue virus 3, Sri Lanka, 2019, Illumina | 10707 | 273 | 27 | 55.05 | 2.58 |
| 13 | KY863456.1, Dengue virus 3 isolate 201610225, Indonesia, 2017, IonTorrent, Sanger dideoxy sequencing | 10707 | 278 | 28 | 52.84 | 2.5 |
| 14 | MH544649.1, Dengue virus 3 isolate 449686_Antioquia_CO_2015, Colombia, 2018, Illumina; Sanger dideoxy sequencing | 10707 | 273 | 28 | 52.84 | 2.49 |
| 15 | MH823209.1, Dengue virus 3 isolate SMD-031, Indonesia, 2019, Illumina | 10707 | 272 | 28 | 52.84 | 2.46 |
| 16 | LC379197.1, Dengue virus 3 strain SYMAV-17/Gabon/2017 genomic RNA, 2019, Illumina | 10641 | 271 | 29 | 52.85 | 2.06 |
| 17 | KY921907.1, Dengue virus 3 isolate SG(EHI)D3/15095Y15, 2017, Singapore, Sanger dideoxy sequencing | 10667 | 266 | 29 | 53.58 | 2.09 |
| 18 | *KF041255.1*, Dengue virus 3 isolate D3/Pakistan/55505/2007, 2013, Sanger dideoxy sequencing | 10675 | 268 | 29 | 53.55 | 2.07 |
| 19 | *LC379196.1*, Dengue virus 3 strain SYMAV-09/Gabon/2016 genomic RNA, 2019, Illumina | 10663 | 273 | 29 | 53.64 | 2.53 |
| 20 | *LC379195.1*, Dengue virus 3 strain SYMAV-07/Gabon/2016 genomic RNA, 2019, Illumina | 10663 | 273 | 29 | 53.64 | 2.53 |
| 21 | *KJ579245.1*, Dengue virus 4 strain DENV-4/MT/BR23_TVP17909/2012, Brazil, 2020, Illumina | 10649 | 273 | 26 | 53.12 | 2.09 |
| 22 | *MG272274.1*, Dengue virus 4 isolate D4/IND/PUNE/IRSHA-FG-03 (S-49), complete genome, India, 2018, Ion Proton System | 10652 | 270 | 27 | 53.07 | 2.07 |
| 23 | *KY672960.1*, Dengue virus 4 isolate DENV-4/China/YN/15DGR394 (2015), 2019, Sanger dideoxy | 10661 | 276 | 26 | 52.63 | 2.09 |
| 24 | *KX224312.2*, Dengue virus 4 isolate SG(EHI)D4/02990Y14, Singapore, 2017, Sanger dideoxy sequencing | 10652 | 275 | 27 | 52.21 | 2.09 |
| 25 | *MG272272.1*, Dengue virus 4 isolate D4/IND/PUNE/IRSHA-FG-01 (1028), India, 2018, Ion Proton System | 10652 | 272 | 27 | 54.57 | 2.09 |

**Table 4.** The Ebola Virus, (GenBank), *atg*-walk

| # | Genbank Virus Name, Registration Year, Sequencing Technology | Nucleotide Number | Number of *atg*-Triplets | Word Median Length | RMS Word Length | Fractal Dimension of the Word-length Distribution |
| --- | --- | --- | --- | --- | --- | --- |
|  | 1 | 2 | 3 | 4 | 5 | 6 |
| 1 | *MG572235.1*, Zaire ebolavirus isolate Ebola virus/H.sapiens-tc/COD/1995/Kikwit-9510621, Zaire, 2019, PacBio; Illumina | 18957 | 329 | 40.5 | 85.29 | 2.03 |
| 2 | *KU174137.1*, Mutant Zaire ebolavirus isolate Ebola virus/H.sapiens-rec/COD/1976/Yambuku-Mayinga-eGFP-BDBV_GP, Zaire, 2019, Illumina | 19774 | 339 | 41.5 | 84.02 | 2.06 |
| 3 | *KY786025.1*, Ebola virus strain Ebola virus/M.fascicularis-wt/GAB/2001/untreated-CCL053D5, Gabon, 2018, IonTorrent | 18871 | 327 | 40.5 | 85.06 | 2.03 |
| 4 | *KY785936.1*, Ebola virus strain Ebola virus/M.fascicularis-wt/GAB/2001/100mg-CA470D5, Gabon, 2018, IonTorrent | 18871 | 327 | 40.5 | 85.06 | 2.03 |
| 5 | *MH121162.1*, Sudan ebolavirus isolate Ebola virus/H.sapiens-tc/Sudan/1976/Boniface-R4142L, 2019, Illumina | 18831 | 345 | 37.5 | 77.79 | 2.2 |
| 6 | *MK952150.1*, Sudan ebolavirus isolate Ebola virus/H.sapiens-wt/SSD/1976/Maridi-BNI/DT, South Sudan, 2020, Illumina | 18847 | 345 | 37 | 77.84 | 2.2 |
| 7 | *MH121169.1*, Sudan ebolavirus isolate Ebola virus/H.sapiens-tc/Sudan/2004/Yambio-HCM/SAV/017, 2019, PacBio; Illumina | 18849 | 345 | 37 | 77.84 | 2.2 |
| 8 | *NC_039345.1*, Bombali ebolavirus isolate Bombali ebolavirus/Mops condylurus/SLE/2016/PREDICT_SLAB000156, Sierra Leone, 2018, Sanger dideoxy, Illumina | 19043 | 325 | 36 | 84.24 | 2.38 |
| 9 | *MW056492.1*, Bombali ebolavirus isolate X030, Kenya, 2020, Illumina | 19025 | 326 | 36 | 84.11 | 2.37 |
| 10 | *MF319186.1*, Bombali ebolavirus isolate Bombali virus/C.pumilus-wt/SLE/2016/Northern Province-PREDICT_SLAB000047, Sierra Leone, 2019, Sanger dideoxy | 19043 | 324 | 36 | 85.27 | 2.36 |
| 11 | *MK340750.1*, Bombali ebolavirus isolate B241, Kenya, 2019, Illumina | 19025 | 328 | 36 | 83.59 | 2.36 |
| 12 | *MW056493.1*, Bombali ebolavirus isolate Z153, Kenya, 2020, Illumina | 19025 | 332 | 36 | 81.46 | 2.41 |
| 13 | *MK028856.1*, Bundibugyo ebolavirus isolate Ebola virus/H.sapiens-tc/Uganda/2007/Bundibugyo-200706291, 2019, PacBio; Illumina | 18940 | 325 | 39 | 84.62 | 2.01 |
| 14 | *MK028834.1*, Bundibugyo ebolavirus isolate Ebola virus/H.sapiens-tc/Uganda/2007/Bundibugyo-R4386L, Uganda, 2019, PacBio; Illumina | 18917 | 325 | 39 | 84.62 | 2.01 |
| 15 | *MK028835.1*, Bundibugyo ebolavirus isolate Ebola virus/H.sapiens-tc/Uganda/2007/Bundibugyo-200706291, 2019, PacBio; Illumina | 18936 | 325 | 39 | 84.63 | 2.01 |

